# Structural Basis of M1 Muscarinic and H3 Histamine Receptor Inhibition in OPC Differentiation

**DOI:** 10.64898/2026.04.01.715893

**Authors:** Bryan Raubenolt, Fabio Cumbo, Jayadev Joshi, William Martin, Satish Medicetty, Yan Yang, Bruce Trapp, Daniel Blankenberg

## Abstract

Muscarinic and histamine receptors are neurotransmitter-binding proteins within the large family of G protein-coupled receptors (GPCRs) and are relevant to human health and disease, including multiple sclerosis (MS), a chronic immune-mediated inflammatory demyelinating disease of the central nervous system (CNS) with neurodegenerative components. MS affects approximately 1 in 333 people, and women are affected at roughly threefold higher rates than men. A major pathological feature of MS is demyelination with incomplete remyelination of axons in the CNS. Because oligodendrocyte progenitor cells (OPCs) can differentiate into mature oligodendrocytes that restore myelin, small molecules that promote OPC differentiation represent a potential therapeutic strategy. High-throughput screening identified 18 hit compounds with EC50 values below 0.2 μM, including the lead compound CN045, which showed an EC50 of 40 nM in vitro. Cheminformatic and experimental target-identification studies implicated the M1 muscarinic receptor and the H3 histamine receptor as candidate targets. To interpret these findings, we performed docking, molecular dynamics simulations, and binding free-energy analyses on complexes involving CN045 and clemastine, a known antihistamine with antimuscarinic activity. The simulations support weaker and less stable binding of CN045 to H3 than to M1 and identify residue-level interactions that contribute to stability within the M1 binding pocket. Comparisons between CN045 and clemastine at M1 further suggest that the two ligands sample different local conformational ensembles, including differences in conserved microswitch behavior associated with active-like versus inactive-like receptor states. Together, these results provide a structural framework for understanding ligand-specific M1 engagement and may help guide future optimization of remyelination-promoting compounds.

## 1 Introduction

Myelin loss on neuronal axons is a major pathological feature of multiple sclerosis (MS) and several other neurodegenerative disorders. Another major driver of MS pathology is autoimmune inflammation. Multiple FDA-approved disease-modifying therapies are available for relapsing forms of MS, but none are approved specifically to promote remyelination [1–6]. An alternative therapeutic strategy is to enhance remyelination and thereby address an important unmet component of MS pathogenesis. At present, no FDA-approved medications are indicated specifically for this purpose.

The biological mechanisms that drive myelin loss (demyelination) in MS involve highly complex cascades of molecular events, including important genomic and transcriptomic contributors [7–12]. Conversely, the biology underlying remyelination is also difficult to quantify. Deciphering these processes is part of a rapidly growing field focused on the treatment and mechanistic understanding of MS and other demyelinating diseases. Oligodendrocytes are glial cells in the central nervous system (CNS) whose primary function is to form myelin sheaths around axons. Oligodendrocyte progenitor cells (OPCs) are the precursors of mature, myelinating oligodendrocytes, and their differentiation provides the cellular basis for myelin formation and remyelination [13,14]. Thus, a new class of therapeutic agents is being explored to promote OPC differentiation directly. In this work, we examine two such compounds to better understand their possible mechanisms of action.

Clemastine is a well-studied compound with antihistamine activity and recognized antimuscarinic activity, including activity at the M1 receptor [15–20], and it is also a strong promoter of OPC differentiation [15–21]. It has been evaluated in multiple clinical studies of remyelination in MS [22–25], with at least one study reporting positive results, particularly for remyelination of the optic nerve in patients with relapsing MS and chronic optic neuropathy [22]. The M1 muscarinic receptor and H3 histamine receptor are transmembrane proteins that belong to the broader family of G protein-coupled receptors (GPCRs). They bind acetylcholine and histamine, respectively, and regulate many essential cellular signaling processes. Although clemastine is classically described as an H1 histamine receptor antagonist/inverse agonist [26], accumulating evidence suggests that it also binds the M1 receptor [27–29]. Disruption of muscarinic signaling appears to favor OPC differentiation, and genetic or pharmacologic inhibition of this pathway has been associated with increased differentiation and remyelination [30,31]. Taken together, the available literature supports muscarinic inhibition as a plausible strategy for remyelination therapy. Clemastine therefore serves as an informative benchmark for the present study and for related drug-development efforts.

Our team conducted high-throughput screening to identify new compounds that promote OPC differentiation. An initial screen of a 20,000-molecule library identified 43 compounds with EC50 values below 1 μM. Of these, 18 had EC50 values below 0.2 μM, and the lead compound showed an EC50 of 40 nM. This publicly available compound (not patented), denoted CN045 (for “Cashel Neural compound 045”), showed notable potency for inducing OPC differentiation in vitro. Figure 1 illustrates the screening workflow. A bar plot is also included to summarize the compounds’ in vitro activities. The vehicle control was dimethyl sulfoxide (DMSO), and benztropine (BenZ), an anticholinergic medication used to treat Parkinson’s disease and other movement disorders, was included as an additional comparator. CN045 produced an approximately twofold increase in activity relative to clemastine, providing a strong rationale for a comparative computational analysis. Understanding how and why this difference arises, through the lens of computational chemistry, is the central focus of the work presented here.

**Fig. 1.**
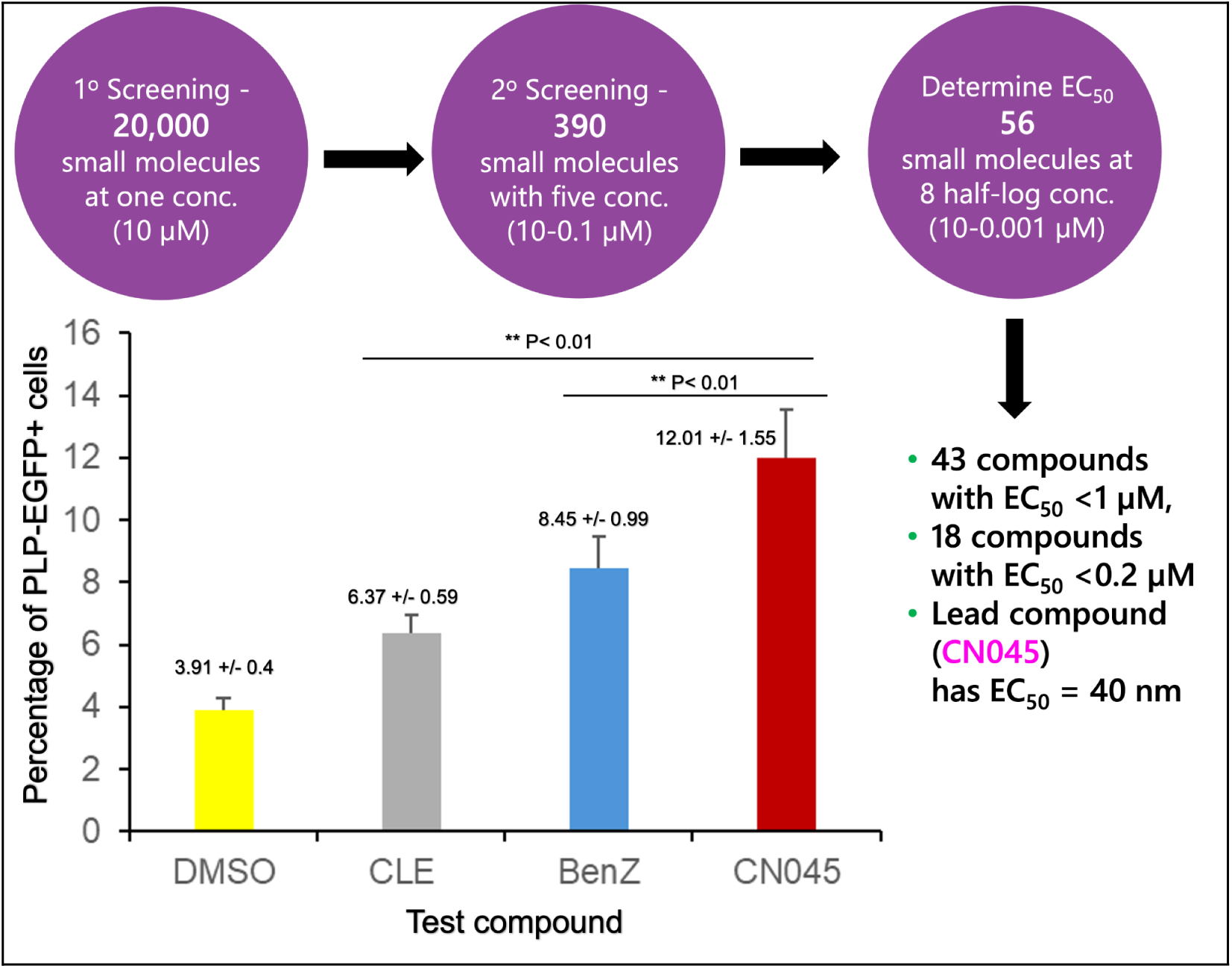
High-throughput screening workflow for OPC-differentiation-inducing compounds. Starting from an initial library of 20,000 molecules, 43 compounds showed submicromolar EC50 values, including 18 with EC50 values below 200 nM (0.2 μM). The lead compound, CN045, showed an EC50 of 40 nM. Its efficacy also stood out relative to the vehicle control (DMSO) and two remyelination-associated, antimuscarinic comparator compounds (clemastine and benztropine). CN045 increased mouse OPC differentiation in vitro. Quantitative analysis showed a significant increase in the percentage of PLP-EGFP+ oligodendrocytes following treatment with CN045, CLE (clemastine), and BenZ (benztropine) relative to DMSO (P < 0.01), with CN045 showing the strongest effect.

While much is known about clemastine’s targets, particularly the M1 receptor, substantial effort was devoted to identifying the relevant target or targets for CN045. Chemoinformatic analyses (conducted by Bristol Myers Squibb) and a CEREP Express Panel screen (conducted by Eurofins) identified the M1 muscarinic receptor (PDB: 5CXV) and the H3 histamine receptor (PDB: 7F61) as candidate targets. Subsequent confirmatory binding assays further indicated that CN045 has greater affinity for M1 than for H3. Through a combination of docking, molecular dynamics simulations, and binding free-energy calculations, our work (1) helps rationalize why CN045 preferentially binds M1 over H3 and (2) compares the binding mechanisms of CN045 and clemastine in complex with the M1 receptor. Collectively, this investigation provides insights that may help define a pharmacophore for OPC differentiation via M1 inhibition and thereby inform future remyelination-focused drug-design efforts.

## 2 Methods

The work presented here is primarily computational and aims to model the biophysical behavior of three protein-ligand complexes of interest: M1 bound to CN045, M1 bound to clemastine, and H3 bound to CN045. To do so, we used a combination of computational chemistry methods, including homology modeling, molecular docking, full protein-membrane system construction and parameterization, molecular dynamics simulations, and multiple trajectory analyses, including binding free-energy estimation (*ΔG_bind_*). Here we discuss the technical implementations of each of these as well as some background information on the methods employed in the associated experimental work.

### 2.1 Initial PDB files and homology modeling

The starting PDB entries for the M1 muscarinic receptor (5CXV [32]) and the H3 histamine receptor (7F61 [33]) were previously solved via X-ray crystallography. As is common for GPCR structures, both models were incomplete. For M1, the missing regions were located primarily at the termini: the first 44 residues of the N-terminus and approximately 27 residues of the C-terminus were absent. Because the functionally and structurally important regions were otherwise present, additional homology modeling of M1 was not required. The initial M1 coordinates were obtained from the SWISS-MODEL Template Library (SMTL) [34], a repository of experimentally determined proteins mostly derived from the Protein Data Bank [35], used for building homology models. The specific template PDB file is 5cxv.1 and can be found here (https://swissmodel.expasy.org/templates/5cxv.1). The H3 receptor was also missing approximately the first 27 N-terminal residues and the last 13 C-terminal residues. In contrast to the M1 structure, which contains an inserted T4 lysozyme/endolysin fusion commonly used in GPCR crystallography, most of the intracellular domain of H3-specifically intracellular loop 3 (ICL3)-was absent. The missing segment spans residues 242-346, corresponding to 105 residues, and interrupts the continuity of the solved structure. Because these regions are highly flexible, the authors of the H3 structure reported that the truncations were necessary to obtain a stable crystallographic construct [33]. At the time of modeling, available SWISS-MODEL entries for H3 showed low confidence in this region, and the published AlphaFold model (https://alphafold.ebi.ac.uk/entry/Q9Y5N1) likewise predicted the intracellular domain with very low confidence and extensive disorder. Although intracellular and extracellular domains can contain ordered elements, a practical solution here was to bypass explicit modeling of most of the H3 intracellular domain and focus primarily on the transmembrane region, which contains the principal ligand-binding site. We therefore used MODELLER [36] as implemented in ChimeraX [37] and used the original PDB file (7F61) as its template. The resulting model bridged the chain break between Gly241 and Arg347 to restore continuity across the truncated ICL3 region. A similar truncation strategy has been used in previous molecular dynamics studies of H3 [38]. This truncation/modeling strategy is illustrated in Figure S10 of the Supplementary Information.

To assign protonation states and estimations of pKa, hydrogens were added to these protein structures using the H++ server [39]. Following this, histidines were carefully inspected and renamed accordingly (HID, HIE, HIP) to ensure they are correctly parameterized by the chosen force fields in downstream steps.

Residues are referenced either by their sequence numbers in the original PDB files (for example, Tyr418 in M1) or by the Ballesteros-Weinstein numbering scheme for class A GPCRs, in which the most conserved position within each transmembrane helix X is designated X.50 and neighboring positions are indexed relative to that reference [40,41]. Under this scheme, Tyr418 corresponds to Tyr^7.53^, meaning that it lies on transmembrane helix 7 (TM7) three positions downstream of the conserved reference residue. Because GPCR helices vary in length, this numbering scheme facilitates structural comparison across receptors and provides a consistent way to identify conserved functional residues.

### 2.2 Molecular Docking

Once the primary PDB files were obtained as described above, a basic alignment procedure was conducted between the new models and the original PDB files, aligning the structures in a manner that minimizes the overall backbone RMSD as much as possible, producing the closest set of coordinates between them. This was primarily done using the RMSD calculator tool in VMD (version 1.9.4) [42]. The VMD package was also used for all visualization and rendered images in this work. While docking was performed on the original crystal structure PDB files, the purpose of this alignment was to maintain the same reference frame in order to subsequently transfer the docked ligands to the final protein models afterwards. This approach is commonly used in practice (especially when starting with incomplete crystal structures), helping retain the docked ligand’s coordinates, geometry, and bonding network before performing the simulations in complex with the protein models. The opposite approach, where the co-crystallized ligands are transferred over to the aligned protein model first, with docking performed using the model instead, could also be employed for an equivalent outcome. For both proteins, the co-crystallized ligand was situated at the respective orthosteric sites. The ligand in M1 is tiotropium (colored in yellow in Figure 2a and b), a well-known inverse agonist/antagonist of the M1 and M4 muscarinic receptors [32]. In H3, the ligand is also an antagonist (colored in black in Figure 2c), referred to as PF-03654746 in the associated work [33]. Given that our binding studies of CN045 and the available literature on clemastine indicate that these compounds likely bind at the orthosteric site, our team leveraged the crystallographic coordinates of the complexes in the original PDB files, giving us a “ground truth” to base our docking efforts on. Thus, the docking grid was created based on these co-crystallized ligands, specifically using RDKit (version 2021.03.5) [43] to create the grid parameters in the original crystal structure PDB files. Both CN045 and clemastine were subsequently docked within this grid using AutoDock Vina (version 1.2.3) [44]. The intermediate steps, including changing file formats for receptors and ligands were largely handled using functionalities in Open Babel (version 3.1.1) [45]. All of these steps were performed and automated in Galaxy [46], using a previous molecular docking tutorial within the Galaxy Training Network as a guiding reference [47,48]. The top 10 binding poses for each ligand were produced and analyzed, using a combination of their rank and pose resemblance to the co-crystallized ligand as a basis for selecting the optimal complex for the final system setup and subsequent molecular dynamics simulations. The comparisons between the docked compounds and the co-crystallized ligands are shown below in Figure 2. In all 3 cases there is a clear overlap between the docked binding poses of the compounds and the co-crystallized ligand’s coordinates, serving as a reasonable starting point for the simulations.

**Fig. 2:**
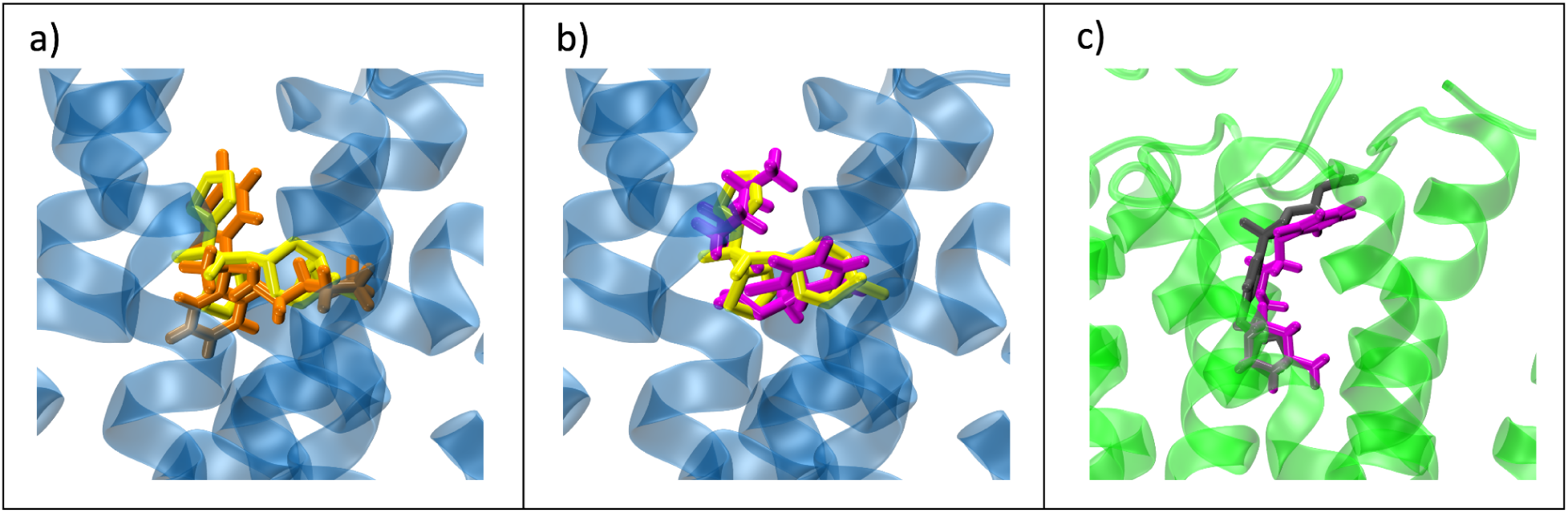
Comparison of the docked poses between a) clemastine (orange) and b) CN045 (magenta) with the co-crystallized ligand (yellow) at the orthosteric site of the M1 receptor (blue), as well as the docked pose of c) CN045 (magenta) with the co-crystallized ligand (gray) at the orthosteric site of the H3 receptor (green). The close proximity between the docking poses and the co-crystallized ligands ensures the simulations start with a reasonably accurate set of coordinates.

### 2.3 Transmembrane system setup and force field application

With the initial protein-ligand complexes defined, the next step was to construct the full simulation systems. Each complex was embedded in a lipid bilayer, solvated with water, and neutralized with ions to achieve a salt concentration of 0.15 M. This was performed in CHARMM-GUI [49], specifically using Membrane Builder and the input generator [50]. While the platform may have been originally developed to work with CHARMM force fields, as the name suggests, support for AMBER force fields was added in recent years [51]. The AMBER FF14SB force field was chosen for the proteins [52], while the ligands were parameterized with the second generation General Amber Force Field (GAFF2) [53] with atomic charges assigned using AM1-BCC [54]. The TIP3P water model [55] was selected to hydrate the system, with the standard Joung-Cheatham parameters used for ions [56]. The lipid bilayer was modeled with Lipid21 [57] using dipalmitoylphosphatidylcholine (DPPC) as a simplified approximation of a neuronal membrane. Figure 3 shows the final systems from three viewpoints. The total atom counts were 80,250 for M1-Clemastine, 70,864 for M1-CN045, and 69,343 for H3-CN045. Differences in atom count arise mainly from modest differences in box size and water content. Full system dimensions are provided in Table S6 of the Supplementary Information.

**Fig. 3:**
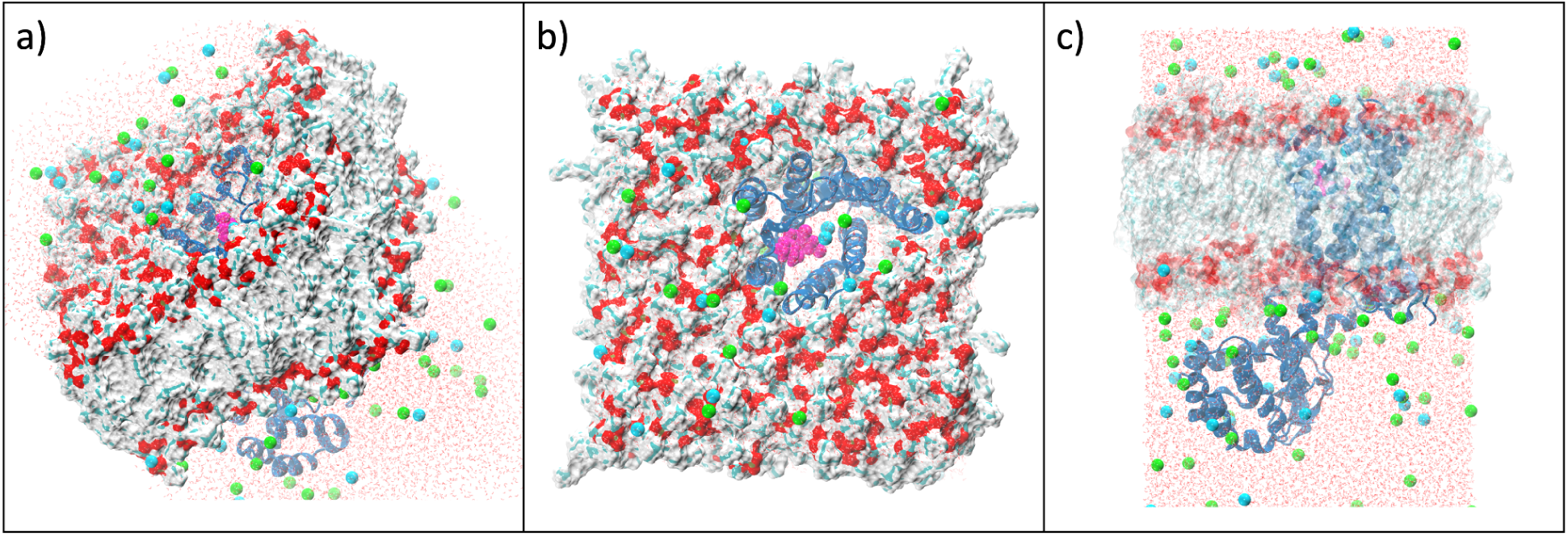
A look at the final transmembrane systems, beginning with a) 3D diagonal view, b) bird’s eye view looking at the extracellular interface (in other words, facing the membrane from the outside of the cell), and c) a slice through the membrane providing a lateral view of the system. Water molecules are depicted as thin red lines, K+ and Cl-ions are spheres colored in green and cyan. The membrane is formed by an upper and lower leaflet of DPPC molecules, with the oxygen atoms in red, hydrogen atoms in light gray, and carbon atoms in light green/aqua. The M1 protein is colored in blue while the CN045 molecule is colored in magenta.

**Fig. 4:**
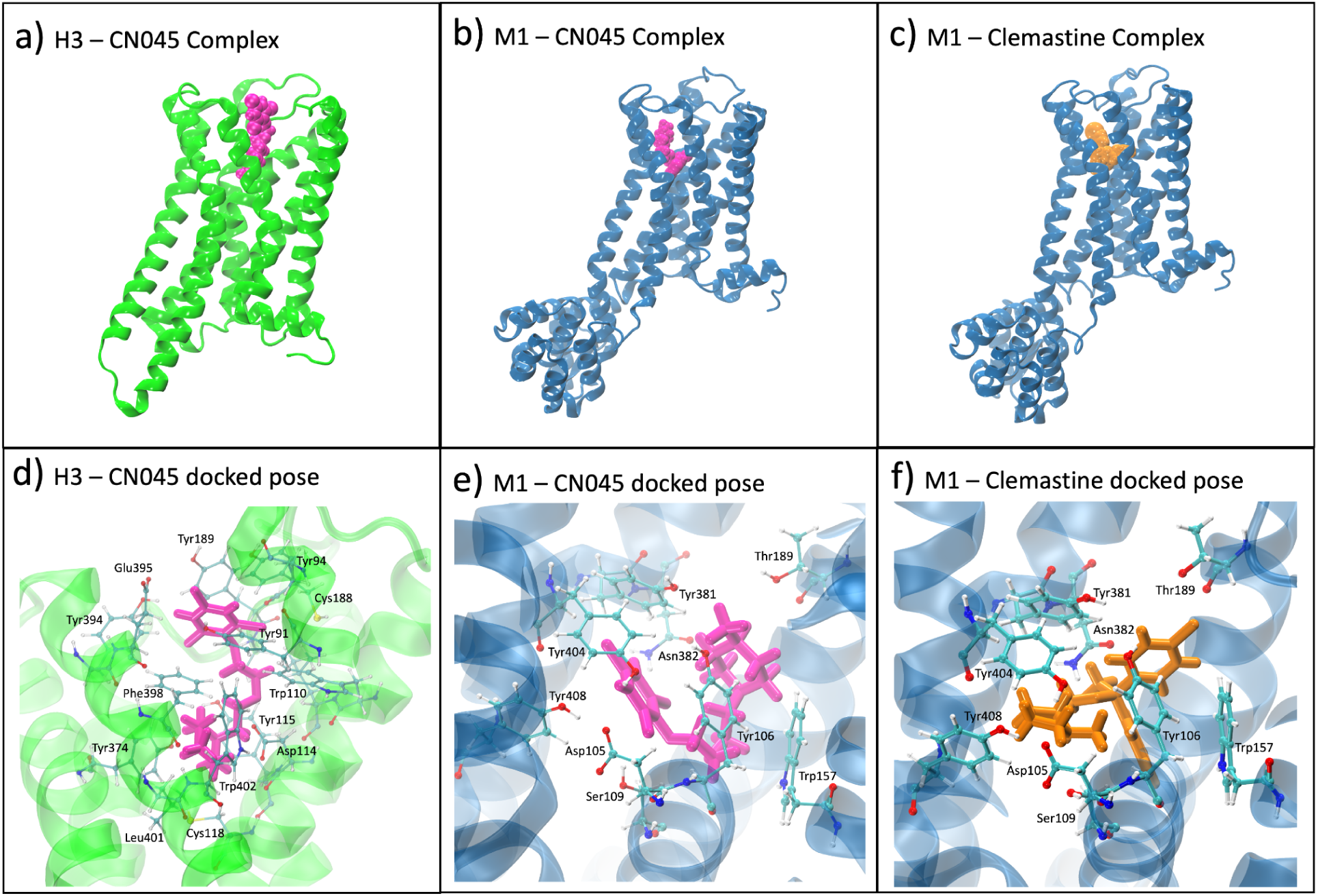
A comparison of individual receptors and the interactions of the docked drug’s conformations with the orthosteric site residues.

### 2.4 Molecular Dynamics Simulations

The initial system setup described above produced GROMACS topology and coordinate files [58], after which a series of energy minimization, equilibration, and production run simulations were performed in order to achieve steady state conditions. Energy minimization was conducted in a series of 5,000 steps with a force tolerance of 1000 kJ/mol/nm, using a steepest descent integrator. The systems were equilibrated at 323.15 K using the Berendsen thermostat [59], through two subsequent equilibrations in the isothermal-isochoric ensemble (otherwise known as the *constant number of particles, volume, and temperature, or NVT, ensemble*), denoted as NVT1 and NVT2. This temperature was chosen to maintain the DPPC bilayer in a fluid phase and should be interpreted as a membrane-modeling choice rather than as physiological temperature. For both equilibrations, 1 femtosecond timesteps were used for a total of 125,000 steps and independent restraints were applied to protein backbones, sidechains, as well as a single lipid molecule serving as an “anchor” for the membrane. Initial atomic velocities were derived from a Maxwell-Boltzmann distribution and assigned in NVT1. The conditions during NVT2 were the same with the exception of generating initial velocities and cutting the restraint forces approximately in half. What followed were 4 additional equilibration steps, this time in the isothermal-isobaric ensemble (otherwise known as the *constant number of particles, pressure, and temperature, or NPT, ensemble*), using the Berendsen barostat for pressure coupling. Each NPT step ran for 250,000 steps and employed a 2 femtosecond timestep, for a total of 1 million steps of NPT equilibration. The primary reason for these discrete sequential runs was to gradually taper off the restraints. Following this, up to 10 short post-equilibration runs were conducted to further equilibrate the system, for a total of 5 million steps, this time using the Parrinello-Rahman barostat [60] and the Nosé-Hoover thermostat [61]. In all cases, 3 groups were used for temperature coupling (namely the solutes, membrane components, and the solvent). The Verlet cutoff scheme with 9 Å cutoffs was employed for van der Waals (*rvdw*) and coulombic interactions (*rcoulomb*), as well as the neighbor list radius (*rlist*). The Particle Mesh Ewald (PME) [62] was used to treat long-range electrostatics, with dispersion corrections applied for energy and pressure. Hydrogen bonds were constrained using the LINCS algorithm [63]. A final, 1 microsecond production simulation was generated, with the central difference in the parameters of these longer simulations being an increase in the cutoffs to 12 Å, as well as the use of a vdw-modifier to handle discontinuity in the Lennard-Jones potential (van der Waals energy) at the cutoff. This method applies a smoothing function that continuously, rather than abruptly, scales the potential energy to zero. To account for uncertainties and variations in the initial velocities, as well as any stochasticity in this protocol, 3 “replica” simulations were run beginning from the minimization step for each system. In doing so, this provides our study with a more realistic, aggregate view of the evolution of these systems. Aside from this final production run, all other simulations (including minimization, equilibrations, and short production runs) used the stock .mdp files (and a bash script to run them) that were included in the GROMACS directory in the downloaded archive produced from CHARMM-GUI’s input generator. These longer replica simulations were all run in Galaxy [46]. Galaxy tools were used to post-process and analyze trajectories, using a combination of underlying tool dependencies including MDAnalysis [64], BRIDGE [65], MDTraj [66], and Bio3D [67], as well as additional analysis functions (including radius of gyration, RMSD, and center of mass distances) using GROMACS 2022 [58]. Another Galaxy tool leveraging plugins from Visual Molecular Dynamics (VMD [42]) was used for the hydrogen bond analyses. With an established history of providing a reliable, fully open-source alternative to proprietary molecular simulation software [68], Galaxy provided our team with an efficient, UI-based option for running and analyzing these simulations, as well as storing and organizing data. The fully automated workflows have also been made publicly available (see Data Availability). VMD was also used to visualize the trajectories and render the images presented in this manuscript.

### 2.5 Binding Free Energy Calculations

As an additional measure of complex stability and comparison between drug candidates, free energy calculations were performed using the Molecular Mechanics Poisson–Boltzmann Surface Area (MM-PBSA) method [69]. This is a popular *end state* free energy calculation method, where the free energy is computed on initial and final system states only (the bound and unbound states in protein-drug complexes, for example), rather than the paths in between. By combining molecular mechanics energies from a force field (rather than more intensive approaches like calculating quantum mechanical energies), along with employing implicit solvation models, it provides computational efficiency while preserving accuracy. The method requires only a single protein-complex simulation trajectory, which makes it even more efficient and another reason for its wide use in drug discovery research efforts [69,70]. For a given protein-ligand system, the binding energy is derived as seen below:

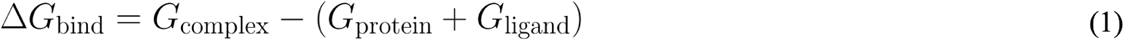

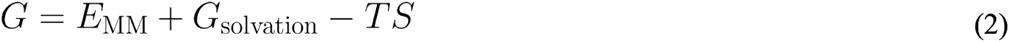

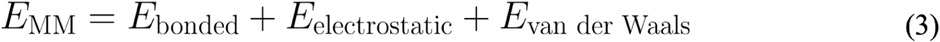

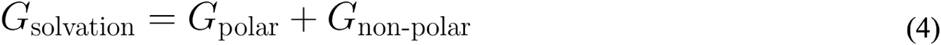

From equation 1, the binding energy described above is defined as the difference between the free energy of the complex (*G*_complex_) and the individual free energies of the protein (*G*_protein_) and ligand (*G*_ligand_). Each one of these free energy terms for each of the system components (complex, protein, and ligand) is defined the same way. They are the sum of the molecular mechanics energies (*E*_MM_) and the solvation energy (*G*_solvation_), minus the entropic contributions (the product of temperature *T* and entropy *S*), as seen in equation 2. *E*_MM_ is made up of the energies governing the standard molecular interactions, namely the bonded interactions within a molecule (*E*_bonded_, otherwise known as the internal energy), the electrostatic or coulombic energies derived from point charges on interacting atoms (*E*_electrostatic_), and the van der Waals energies (*E*_van der Waals_), which encapsulate the effects of induced dipole moments, among other properties. Each one of these energy terms is distance-dependent and is derived from a specified force field (usually the same force field used to generate the trajectories to begin with).

Depending on the systems being studied, the entropic contributions can be neglected in practice, due to the high computational cost and sensitivity to noise. For comparative studies involving the same receptor and structurally similar ligands, these terms often lead to negligible differences in the final *ΔG_bind_*, and thus may be considered constant and omitted. Since the MM-PBSA method can be seen as more of a semiquantitative ranking tool rather than a rigorous alchemical free-energy method (such as thermodynamic integration or free energy perturbations), neglecting the entropic contributions in these scenarios is a reasonable approximation that still preserves relative scoring and provides a comparison of thermodynamic stability between different protein-ligand complexes.

The remaining component of the calculation is the solvation energy (*G*_solvation_), which is composed of both polar (*G*_polar_) and non-polar (*G*_non-polar_) contributions, as seen in equation 4.

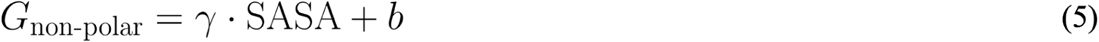

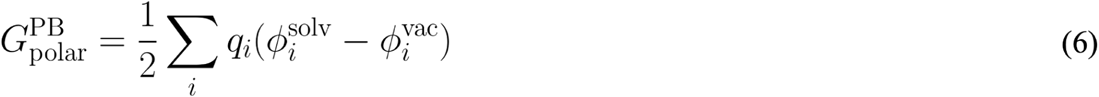

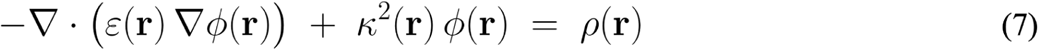

There are two forms of computing *ΔG_bind_*, namely the Generalized Born (MM-GBSA) and the Poisson–Boltzmann (MM-PBSA) methods. Both approaches are the same up until equation 5, the non-polar solvation energy. Here, *γ* is the surface tension coefficient, which is intended to represent the cost of exposing a hydrophobic surface to water. The other parameter, *b*, is a constant which approximates the cavitation energy (the cost of forming a solute-shaped cavity in water) and the dispersion interactions between solute and solvent. Both of these are empirically derived parameters, usually determined from a linear regression fitted to experimentally determined hydration free energies. The remaining term is the solvent accessible surface area (SASA). The efficiency of these methods is aided by the inherent use of implicit solvent models. Instead of using explicit solvent molecules, approximations are made to capture these effects, including the use of SASA in equation 5, which makes the non-polar term a function of the surface area that an explicit number of water molecules would theoretically be exposed to.

The core difference is how the polar solvation energy (*G*_polar_) is calculated. While both methods employ an implicit solvent approach here as well, by treating the solvent as a continuous dielectric medium, there are key differences in how this is done. Equation 6 is the equation for *G*_polar_ in MM-PBSA calculations. It is the sum over all atomic reaction fields, computed as each partial charge *q_i_* multiplied by the difference between the electrostatic potential in solvent (*Φ^solv^*) and in vacuum (*Φ^vac^*) at that atom’s position. These potentials are determined by solving the PB equation (7). The terms on the left are comprised of a gradient operator ⛛ (a vector representing the potential’s changes in all 3 cartesian directions), the spatially varying dielectric constant દ*(r)*, the electrostatic potential *Φ(r)*, and the inverse Debye screening length *k(r)*, which quantifies how strongly mobile ions in the solvent screen or damp the electrostatic potential (effectively capturing the influence of salt concentrations in the solvent). The term on the right, *ρ(r)*, is the molecular charge density, which is represented as a sum of Dirac delta functions centered at each atomic position, each weighted by that atom’s partial charge. The PB equation is solved twice on the same spatial grid but with different dielectric maps for દ*(r)*: 1) once with the true heterogeneous dielectric, where દ*(r)* is low (≈ 1–4) inside the solute cavity (દ*_in_*) and high (≈ 80) in the surrounding solvent (દ*_out_*), to obtain the solvent-screened potential *Φ^solv^*. 2) once with a uniform low dielectric (દ*(r)* = દ*_in_* everywhere) and no ions, to obtain the vacuum potential *Φ^vac^*. These potentials are then used to determine *G*_polar_. Equation 7 is known as the linearized version of the PB equation, which retains the effect of mobile ions but approximates their response as linearly dependent on the electrostatic potential, rather than exponentially.

MM-PBSA is often more physically realistic than MM-GBSA for charged or solvent-exposed systems because it can better represent dielectric boundaries and electrostatic screening, albeit at a modestly higher computational cost. This increased cost arises because a partial differential equation must be solved numerically to capture these effects (7), rather than relying on the analytical approximation used in the generalized-Born formalism. Accordingly, while MM-GBSA remains useful for rapid relative scoring, MM-PBSA can provide a more detailed estimate of the polar solvation term and may improve comparative binding-energy estimates *ΔG_bind_*in systems such as those studied here.

Each replica trajectory was made up of 5000 frames corresponding to 1 µs of simulation time. To help ensure steady state conditions, only the last 100 frames (representing the last 20 ns) were extracted for these calculations. The exception to this was with replica 1 of the H3-CN045 system, where frames 700-800 (corresponding to approximately 140-160 ns) were used instead. This was done because CN045 left the binding pocket early in the simulation, which would have yielded incomparable binding energies with respect to the other replicas. The selected trajectory window spans the period in which the complex remains initially stable and then begins transitioning toward a less stable, more dynamic interaction pattern. This ensured that the MM-PBSA calculations were performed on frames where the complex was still formed, but also representative of its eventual behavior.

These calculations were carried out using gmx_mmpbsa [71]. This package is based on the original Python-wrapped AmberTools version [72], and introduces several enhancements. Most notably, it was developed to work directly with GROMACS files, including trajectories, topology, coordinate, and index files. The tool also provides a useful GUI for a quick analysis of the data. The final plots used in our analysis, however, were generated using in-house Python scripts (which have been made available to the public; see Data Availability), giving our team greater flexibility in visualizing the full breadth of data from the MM-PBSA calculations.

### Statistical comparison of binding free energies

The final binding free energy estimates from three independent molecular dynamics replicas were summarized for each system using two approaches: an unweighted analysis (nonparametric Monte Carlo) and a weighted analysis (parametric Monte Carlo).

In the unweighted analysis, each replica mean was treated as one equally weighted observation. We performed nonparametric Monte Carlo resampling by drawing (with replacement) from the set of replica means for each system, pairing these draws, and estimating the probabilities P(A < B) as the fraction of draws where system A was lower (more negative) than B.

In the weighted analysis, per-replica variability (SD(Prop.)) was used to construct inverse-variance weights, such that replicas with lower variability contributed more strongly to the combined estimate. Because molecular dynamics trajectories exhibit temporal correlation, these variability estimates may not correspond to strictly independent sampling of the replica mean. Accordingly, this weighted analysis is presented as an exploratory, heuristic sensitivity analysis to assess how differences in replica variability influence the inferred probability that one system exhibits a more favorable binding free energy. Inverse-variance weights *w_i_* were calculated, and a weighted mean and corresponding uncertainty (SE*_w_*) were obtained:

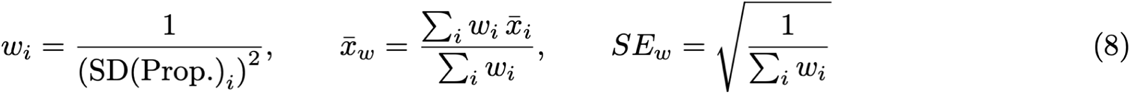

Parametric Monte Carlo sampling was then performed by drawing from the normal distribution N(x̄_w_, SE*_w_*) for each system, independently sampling from each system and comparing paired draws, to compute P(A < B) as in the unweighted analysis. These probabilities represent the Monte Carlo–estimated likelihood that one system exhibits a more favorable (lower) binding free energy than the other, conditional on the observed replica means and their variability.

### 2.6 Experimental data

#### 2.6.1 Binding assays

Binding assays were performed by Eurofins (https://www.eurofinsdiscovery.com/), using the *CEREP Express Panel*, which provided inhibition strengths at a fixed concentration, for 55 different protein targets (see Figure S4). In each one of these experiments, compound binding was quantified as a percent inhibition of the binding of a reference compound, a radiolabeled ligand known to specifically bind to the receptors. These reference compounds were *pirenzepine* for the M1 muscarinic receptor and *(R)*α*-Me-histamine* for the H3 histamine receptor.

The results in the plots seen in Figures S2-S4 as well as Tables S3 and S4, are expressed as a percent of control specific binding (of the reference compound):

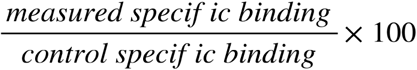

as well as the percent inhibition of control specific binding obtained in the presence of the test compounds (CN045 and clemastine):

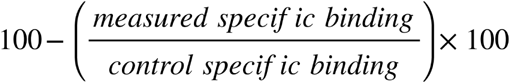

The IC_50_ values correspond to the test compound’s concentration causing a half-maximal (at least 50%) inhibition of control specific binding. These values and Hill coefficients (nH) were determined through a nonlinear regression analysis of the competition curves generated with mean replicate values by fitting the curves to the Hill equation:

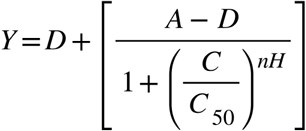

where Y = specific binding, A = left asymptote of the curve, D = right asymptote of the curve, C = compound concentration, C_50_ = IC_50_, and nH = slope factor.

The inhibition constants (K_i_) were calculated using the Cheng-Prusoff equation:

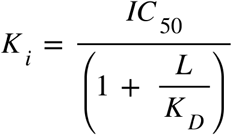

where L = concentration of radioligand in the assay, and K_D_ = affinity of the radioligand for the receptor. A Scatchard plot was used to determine K_D_.

Software developed at Cerep (Hill software) was used for this analysis and validated by comparison with data generated by the commercial software SigmaPlot® 4.0 for Windows® (© 1997 by SPSS Inc.).

#### 2.6.2 Primary mouse oligodendrocyte progenitor cell preparations

Primary mouse OPCs were obtained from neurospheres generated from embryonic day 14.5 brains of PLP-EGFP transgenic mice as described previously. Dissociated single-cell suspensions were plated on poly-L-ornithine-coated 96-well plates (Sigma) at a density of 6,000 cells per well. After two days of proliferation, OPC cultures were treated with clemastine, benztropine, or CN045 for 5 days in differentiation medium. DMSO served as the negative control. After 5 days, cells were fixed in 4% PFA, stained with Hoechst to label nuclei, and imaged. The percentage of PLP-EGFP-positive cells was determined using a Cytation 5 Cell Reader (Agilent BioTek). Twenty images were acquired per condition, and image-analysis algorithms were used to quantify total PLP-EGFP+ oligodendrocytes, total nuclei, and pyknotic nuclei.

## 3 Results

The main goals of these simulations were to provide a structural and thermodynamic basis for (a) why CN045 binds more strongly to the M1 muscarinic receptor than to the H3 histamine receptor and (b) how CN045 and clemastine, both strong inducers of OPC differentiation, differ when bound to M1. For each replica, we evaluated structural descriptors (RMSF, RMSD, and R_g_) for both receptor and ligand, together with binding free-energy calculations and geometry-based hydrogen-bond-like contact analyses. Collectively, these data provide insight into the determinants of complex stability.

### 3.1 Cheminformatics analysis of the compounds

The tested compounds share notable physicochemical properties, which further help explain their observed biological activity and binding. Figure 5 below highlights some of these properties, in particular those most relevant to Lipinski’s Rule of 5 [73]. These calculations were performed using the SwissADME server [74]. The full ADME analysis is included in Table S1 of the Supplementary Information, as well as a 3D representation of the distribution of these metrics in Figure S1.

**Fig. 5:**
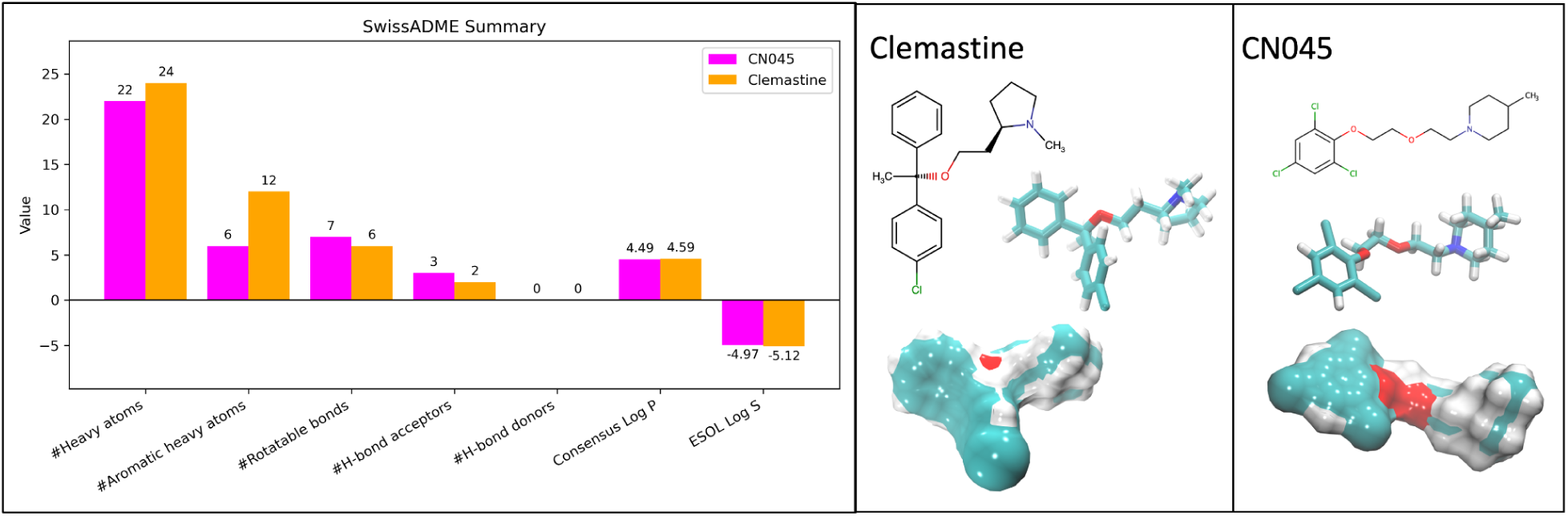
Comparison of physicochemical properties and molecular structures of both compounds.

Both compounds are largely hydrophobic and contain at least one aromatic group and one methyl-substituted heterocyclic amine connected by an aliphatic ether chain. Several differences distinguish them. CN045 contains a trisubstituted aryl halide, whereas clemastine contains a monosubstituted aryl halide; in both cases, chlorine is present. The ether linker is longer in CN045, spanning six heavy atoms rather than three as in clemastine, and CN045 contains two ether oxygens whereas clemastine contains one. The terminal heterocyclic amine also differs: CN045 contains a substituted piperidine ring, whereas clemastine contains a substituted pyrrolidine ring. These differences are apparent in Figure 5, which shows both 2D chemical structures and 3D licorice-and-surface representations.

The bar plot and table further highlight the strong similarity between the compounds. Their molecular weights differ by less than 23 g/mol, and their heavy-atom counts are also similar (22 versus 24). The number of rotatable bonds is likewise close (7 versus 6), indicating comparable overall flexibility, although CN045 may be somewhat more conformationally adaptable because of its longer linker and lack of a second aromatic ring. Neither compound contains classical hydrogen-bond donors, whereas both contain acceptors, including ether oxygens and tertiary amine nitrogens. Because CN045 contains an additional ether oxygen, it has a somewhat greater hydrogen-bond accepting capacity in the neutral descriptor representation. Both compounds also show very similar lipophilicity (LogP) and solubility (LogS), consistent with high predicted GI absorption. Both satisfy Lipinski’s rule-of-five criteria, supporting their general drug-like character. Figure 5 also helps visualize why both molecules may engage overlapping targets involved in OPC differentiation.

### 3.2 Binding assays

The initial screening using the CEREP Express Panel (conducted by Eurofins) derived inhibition potencies for CN045 against 55 protein targets, including muscarinic receptors, histamine receptors, gamma-aminobutyric acid receptor (GABAr), ion channels, and many others in the GPCR family. In all cases, the same concentration was used for CN045 (3.0E-06 M). The results of this screening are presented in Figure S4. From this data, it became clear that CN045 binds the strongest to the M1 muscarinic receptor, with an 81.7% inhibition rate at the tested concentration, followed by 73.5% with the M2 muscarinic receptor and 51.1% with the Na^+^ ion channel. Binding to the M3 receptor was notably weaker, at 25.5% inhibition. This suggests stronger engagement of M1 and M2 compared to the other muscarinic subtypes tested in the single-concentration screen.

Separately, a cheminformatics analysis was performed on CN045 and 64 additional compounds from the in vitro OPC-differentiation study. This analysis was conceptually the inverse of a docking-style screen: protein targets were prioritized based on chemical similarity between the test compounds and co-crystallized ligands, together with complementarity between compound chemistry and target binding sites. The results are shown in Figure S5. Targets were ranked by the number of test compounds matched to each receptor. Among the top 25 targets, approximately 84% were GPCRs. The top two predicted targets were H3 and GABAr. Taken together with the CEREP screen, these results motivated follow-up binding assays for CN045 against H3, GABAr, and M1.

The follow-up binding assays are shown in Figure S2 (with the exception of GABAr, which showed virtually no inhibition). M1 and H3 were tested at five concentrations of CN045 ranging from 1.00E-07 M to 1.00E-05 M (Table S3). The IC50 for M1 was approximately one order of magnitude lower than that for H3 (about 1 μM versus greater than 10 μM), further supporting M1 as the more likely target.

With attention now focused on M1, additional experiments were performed to compare CN045 with clemastine at this receptor. The resulting binding assays are shown in Figure S3. Both ligands were tested at 10 concentrations ranging from 3.91E-08 M to 2.00E-05 M (Table S4). A clear difference in IC50 was observed between the two compounds. In this assay, CN045 yielded an IC50 of 1.5 μM, whereas clemastine showed substantially greater apparent potency, with an IC50 below 3.9E-08 M. As shown in Table S4, approximately 85% inhibition was already observed at the lowest tested concentration for clemastine.

### 3.3 Structural analysis of the compounds within the complex

The subtle differences in the physicochemical properties described above can ultimately lead to variations in biological outcomes, as well as the molecular simulations that are meant to represent these conditions. Even when the same force field is used, differences in atom types and charge distributions lead to different potential energy surfaces and interaction patterns. As a result, two simulations that begin from comparable poses in the same binding site will still diverge over time. Thus, there are key structural details that one can monitor in these simulations that provide useful insight into the overall stability of the complex and subsequently correlate to the observed *in vitro* binding affinities. For this, we measured center-of-mass distances (COMd), root mean square deviation (RMSD), and radius of gyration (R_g_), which together give us an idea of residence times within the pocket, as well as conformational stability of the compound.

#### 3.3.1 Comparing the M1-CN045 and H3-CN045 complexes

The binding assays showed that the IC50 for CN045 was about an order of magnitude higher at H3 than at M1. In other words, substantially less CN045 was required to inhibit binding at M1 than at H3. These results suggested stronger binding to M1 and therefore motivated the expectation that the M1-CN045 complex would be more stable in simulation. Part of the work presented here was therefore to test that hypothesis using molecular dynamics metrics designed to evaluate complex stability.

Figure 6 includes the plots measuring COMd between the compounds and the receptor’s orthosteric residues (top row), as well as RMSD (middle row) and R_g_ (bottom row) of the compounds, for each system for the entire 1 microsecond simulation. The replica simulations of each system are colored in black, blue, and red. When comparing the COMd, we see that in M1, for all 3 replicas with CN045 bound, the compound remains very close to its initial coordinates, maintaining a consistent average COMd of approximately 2.1-2.3 Å. In H3, a similar observation is made in replicas 2 and 3. However, in replica 1, there is an obvious instability that arises early on in the simulation. Beginning around 100 ns, CN045 begins to drift away from the H3 binding pocket, reaching a COMd of approximately 3.0 Å by 250 ns. At this point, CN045 leaves the binding pocket entirely for the rest of the simulation. This indicates that CN045 could have a propensity to unbind from H3 more easily, as observed in at least ⅓ of the simulations. This is clearly not the case with any of the M1 replicas, as CN045 remained bound throughout the entire simulation.

**Fig. 6:**
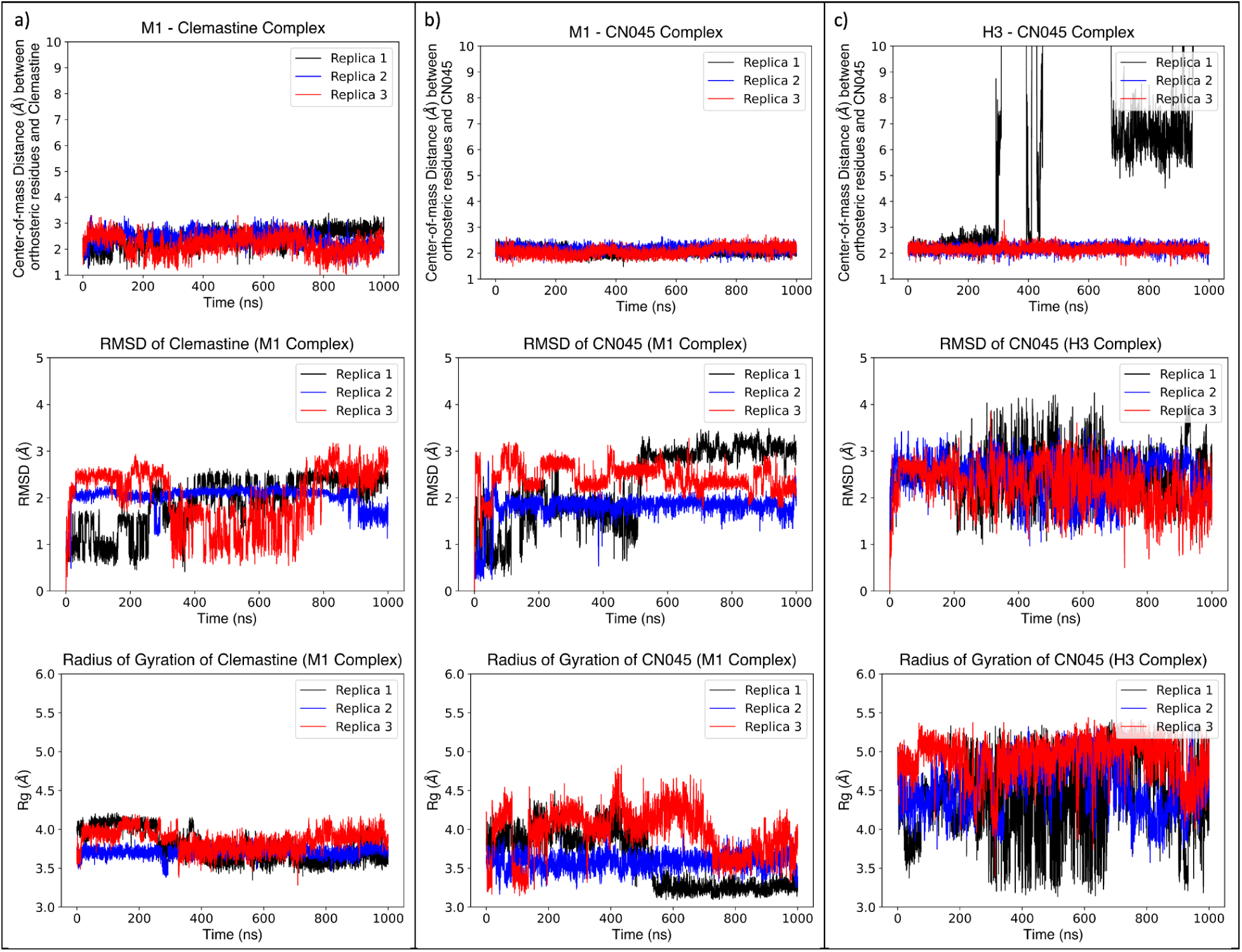
Structural analysis of the binding compounds in each system, throughout 1 microsecond of post-equilibration sampling time. From right to left the data corresponds to the M1-Clemastine, M1-CN045, and H3-CN045 complexes, respectively. From top to bottom, center-of-mass distances are measured between each compound and the orthosteric site residues, as well as the RMSD and radius of gyration (Rg) of the compounds, respectively. All data is measured in angstroms (Å).

The RMSD was measured to monitor how the overall shape (conformation) of the compounds is changing over time (middle row of Figure 6). In M1, we see that CN045 conformations vary slightly among the replicas. In replica 1, there is a consistent conformation observed for the first 500 ns or so (measured with an RMSD of 1.75 Å). At this point the conformation shifts slightly to an RMSD of 3.0 Å or so, consistently remaining here for the remainder of the simulation. In replica 3, we see the predominant conformation hovering around 2.5 Å throughout the simulation. While there are alternating conformations observed with shifts between 2.8 and 2.2 Å, there is little variance observed within these transient conformations. In other words, while the shapes might shift a little more often in replica 2, each of these shapes remain relatively rigid for the respective time periods. Replica 2 appeared to produce the most stable trajectory for CN045, with a relatively consistent RMSD close to 1.8 Å throughout the entire simulation. When compared to the H3 simulation, the difference in CN045’s conformational behavior is somewhat obvious. Across all 3 replicas, there appears to be a significant variance per frame throughout the entire trajectory. This seems most pronounced for replica 1, which would make sense given that CN045 was ejected from the binding pocket and spent most of the simulation diffusing in the bulk solvent/membrane environment. As seen from the COMd plot, however, this was not the case with replicas 2 and 3. This essentially means that, while CN045 remained in the orthosteric site for the entire simulation in these replicas, the molecule was more unstable, rapidly changing conformations between frames. This more erratic fluctuation was not observed when bound to M1, further emphasizing a greater instability in the H3-CN045 complex.

The R_g_ was also used to keep track of major conformational changes in CN045 (bottom row of Figure 6). This is a measure of how compact or extended a molecular structure is, relative to its center of mass. A lower R_g_ implies a more tightly compact conformation, while a higher value corresponds to a more extended conformation. In replica 1 of the M1-CN045 system, we observe a similar transition around 500 ns as we did for the RMSD. The R_g_ here reduces to roughly 3.2 Å, forming a more compact structure, while the RMSD increases in the same time interval (seen in the middle row above). While these two metrics are not identical, when changes are observed in both at the same point in the simulation it helps further emphasize how the structure is changing. This is not always the case however, as can be seen with replica 2, where there is a notable reduction in R_g_ around 700 ns, but there isn’t a significant change in RMSD during that time. In replica 3, the R_g_ remains rather constant (at roughly 3.5 Å) throughout the simulation, which also coincides with the stable RMSD. When compared to the H3 simulations, a similar pattern is once again observed, where there seems to be more fluctuations and variance in R_g_ (as was the case with RMSD). This is most obvious in replica 1, since the compound is outside the binding pocket most of the simulation, but this still holds true in replicas 2 and 3. Furthermore, among replicas 2 and 3, the R_g_ was measured at or above 4.5 Å for the majority of the simulation, while mostly remaining below 4.0 Å in the M1 complex. This indicates that on average, CN045 occupies a more extended conformation when bound to H3, and a more compact conformation when bound to M1. This is caused by a combination of the docked coordinates already reflecting this (Figure 2), as well as the interactions with the binding residues once the simulations began.

In summary, CN045 appears to be less stable when bound to H3 as opposed to M1. This is demonstrated by the compound leaving the orthosteric binding site entirely in at least one H3-CN045 simulation, as well as the increased variance in RMSD and R_g_ throughout the replicas, when compared to the M1-CN045 simulations.

#### 3.3.2 Comparing the M1-Clemastine and M1-CN045 complexes

The same structural analysis was conducted for clemastine in the M1-Clemastine complex. As opposed to the previous analysis, in this case the compound’s behavior appears to be more comparable to what is observed in the M1-CN045 complex, as seen in Figure 6. With respect to COMd, clemastine remains within the binding pocket during all 3 replicas. However, the average distance appears to be closer to 2.5 Å, rather than 2.0 Å (as seen with CN045). This could be due, in part, to the difference in COM of both molecules (clemastine is a bulkier compound with a shorter linker chain and an extra aromatic ring, for instance). Minor fluctuations were noticed in replica 3, but mostly within a relatively small threshold.

For replica 1, two predominant conformations are noted. For the first 250 ns, clemastine demonstrates an RMSD of around 1.0 Å, with a gradual rise to 2.0 Å until 400 ns. The RMSD remains relatively constant throughout the remainder of the simulation, implying a consistent conformation. This transition is similar to what is seen in replica 1 of the M1-CN045 complex, in that case occurring around 500 ns. In replica 3, two predominant conformations are also observed. The first occurs between 0 and 350 ns, corresponding to an RMSD of 2.5 Å. The second conformation takes place between approximately 375 and 775 ns, with a 1.5 Å RMSD. For the remainder of the simulation, the RMSD rises to 2.5 Å, likely reverting back to the previous conformation. As is the case with CN045, clemastine maintains a rather constant conformation in replica 2, corresponding to an RMSD of 2.0 Å for nearly 90% of the simulation.

The R_g_ plots are consistent with the RMSD observations presented above. In replica 1, R_g_ decreases around 250 ns, from roughly 4.1 to 3.6 Å, and remains here for the remainder of the simulation. This coincides with the increase in RMSD at the same point in the simulation. In replica 2, R_g_ remains constant at roughly 3.7 Å for the entire simulation, also consistent with the constant RMSD. In replica 3, we observe the first transition around 375 ns, with R_g_ reducing from 4.0 to 3.7 Å, and increasing once again to 4.0 Å around 775 ns. This also coincides with the predominant RMSD changes in this replica.

All things considered, the conformational dynamics of CN045 and clemastine, when bound to M1, appear to be somewhat comparable. A notable exception could be the greater degree of fluctuation in both RMSD and R_g_ in replica 3. This could be attributable to many factors, including the additional rotatable bond and presence of only one aromatic group in CN045, which inherently increases flexibility in the molecule. Images of the predominant conformations discussed here can be seen in Figure S9.

### 3.4 Structural analysis of the orthosteric binding site

Another important structural component of the system requiring a thorough analysis are the orthosteric-site residues, since these are the primary residues interacting with the compounds in the simulations, and likely *in vitro*. A common metric used, particularly for understanding the effects a binding partner has on a protein, is the root mean square fluctuation, or RMSF. This fundamentally measures the degree of flexibility in a series of residues. Higher RMSFs imply greater flexibility (the residues are moving around a lot more), while lower RMSFs mean that they are more rigid. Figure 7 below shows the measured RMSFs across all systems, with and without the compounds.

**Fig. 7:**
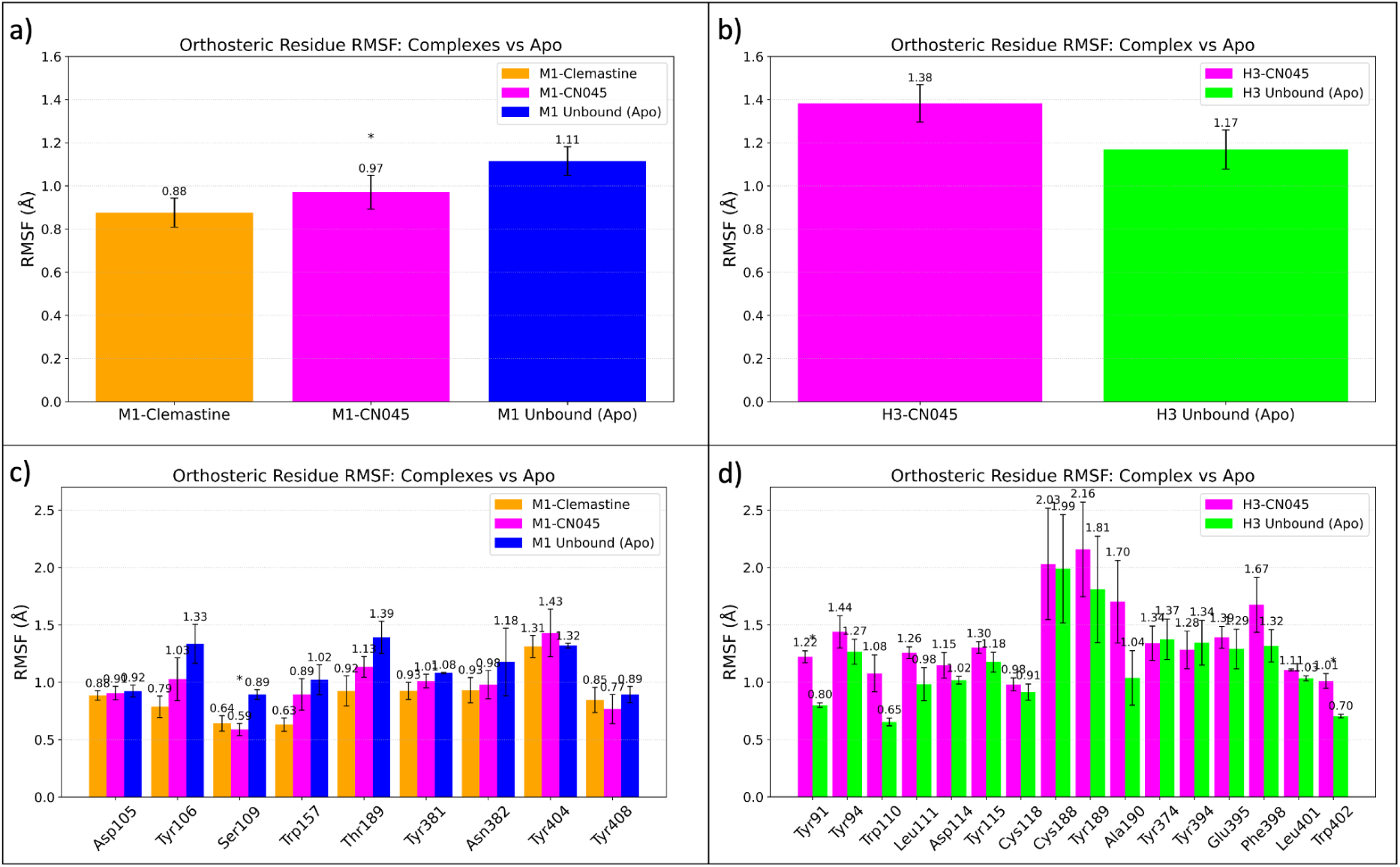
Overall RMSF analysis of the orthosteric residues averaged across the complex (a, b) as well as per residue RMSFs (c, d) for the M1 complexes and H3 complex. The most statistically significant differences (based on p-values less than or equal to 0.05) are highlighted with asterisks. These values correspond to the RMSFs measured at each residue’s alpha carbon.

Generally speaking, in the context of protein-ligand binding, stable complexes lead to lower RMSFs in the binding residues, when compared to the unbound (ligand-free or “apo”) protein. This intuitively means that the interactions with the ligand are strong enough to cause a sort of “stiffening” effect in these residues, which can be quantified. On the other hand, if there is an increase in the RMSF when a ligand is bound, this implies that the complex is less stable, and the binding residues are fluctuating more rapidly. Thus, when comparing different protein-ligand systems, this can further inform which ligand (the drugs in this case) is a more stable binding partner.

Figure 7a and 7c provide the RMSF comparisons between the M1-Clemastine (orange), M1-CN045 (magenta) complex, and the M1 receptor in the unbound state (blue). Both sets of plots are averaged across the replicas of each system. In Figure 7a, we observe the average across all orthosteric site residues. The pattern here indicates that the residues have the lowest RMSF when bound to clemastine (0.88 Å), followed by CN045 (0.97 Å), and the highest in the unbound state (1.11 Å). On the other hand, Figure 7c the RMSF is presented on a per-residue basis. By and large, in both systems, most residues have a reduced RMSF in the bound state. The most pronounced cases include Tyr106, Ser109, Trp157, Thr189, and Asn382. For the remainder of the residues, RMSFs comparable to the unbound state were observed, with only one residue showing higher RMSFs in the complex, Tyr404. Thus, the compounds appear to stabilize most of the orthosteric residues in the M1 receptor.

In Figure 7b and 7d, we provide the RMSFs of the H3-CN045 complex (magenta) and the H3 receptor in the unbound state (green). Contrary to what was observed in the M1 complexes, here we see that the RMSF increases in the bound state. This means that in the unbound state, on average, the orthosteric binding site is more stable. Looking at the corresponding per residue RMSF (Figure 7d), we see that nearly across the board, the RMSF is increasing in the bound state. The most obvious cases are observed with Tyr91, Trp110, Tyr189, Ala190, and Phe398. There were only a couple of residues where the RMSF was marginally less than in the unbound state, with Tyr374 and Tyr394. It is worth mentioning that since CN045 leaves the binding pocket after the first 300 nanoseconds in replica 1, these residues would have now been in the unbound state for the remainder of that simulation. So, the aggregate RMSF of the orthosteric residues in replica 1 in this case is directly influenced by CN045 binding approximately only 30% of the time. On the other hand, the other two replicas had a 100% residence time of CN045, meaning that the RMSF is entirely influenced by interactions with CN045. Overall, the data here therefore indicates that the H3-CN045 complex is relatively unstable, which is consistent with both the somewhat erratic structural behavior of CN045 in Figure 6c and the weaker *in vitro* binding data in Figure S2.

### 3.5 Bonding analysis

There are several forces that contribute to protein-ligand binding, with a significant contributor being the formation of hydrogen bonds. The left column of Figure 8 below illustrates the sorted list of the top 20 drug-residue hydrogen-bond-like pairs by % occupancy, with the y-axis labels specifying the direction of relationship (donors on the left of the arrow, acceptors on the right). The number of replicas for which each pair was observed is written in parentheses (for example, when n=3, that pair was present in all 3 replica simulations of that system). The average number of bonds per frame, across the replicas of each system, is shown in the right side column of Figure 8. When comparing the M1 complexes, one of the first observations is that the number of hydrogen-bond-like contacts maintained throughout the simulations is nearly identical – the mean of the moving average is 15.64 vs 15.58 for the clemastine and CN045 complexes, respectively. The subtle differences between these can be highlighted when observing the % occupancy plots.

**Fig. 8:**
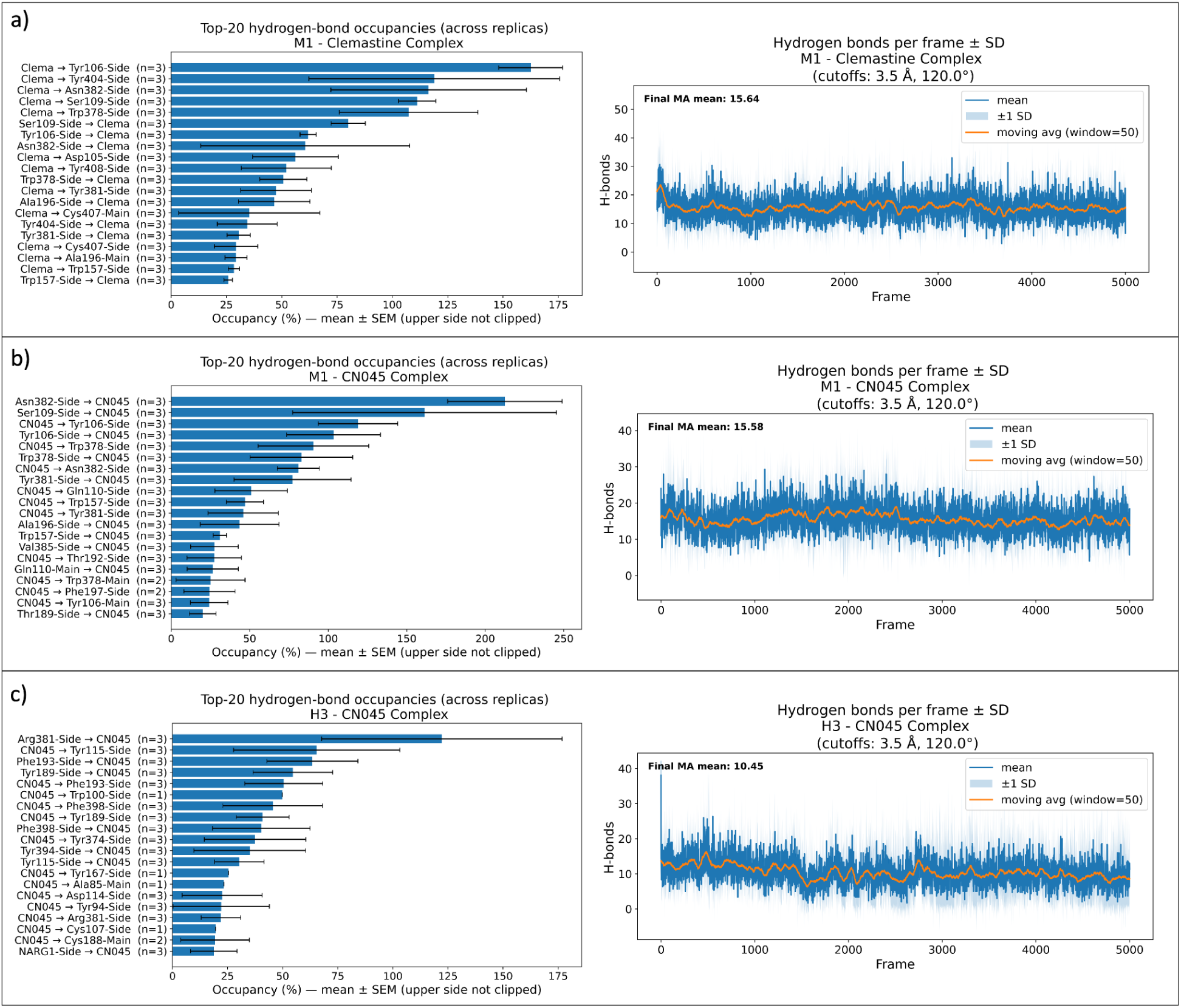
Bonding analysis of all 3 protein-ligand complexes.

These occupancy plots are sorted based on highest to lowest % occupancy throughout the simulation. Many of these plots go beyond 100% occupancy. Although this might seem counterintuitive, the reason is straightforward – for each drug-residue pair, there could be more than one hydrogen-bond-like contact formed depending on the residue. Clemastine, for example, has 2 hydrogen bond acceptors (see Figure 5). Considering the default cutoff criteria employed in the calculation, namely a 3.5 Å distance and a 120° 3-body angle between the donor-hydrogen-acceptor atoms, there would be instances in the trajectories where a single donor is forming a hydrogen bond with each acceptor group on clemastine. If one hydrogen bond is within these cutoffs for 100% of the frames in the simulation, and the other is only established 50% of the time, then total occupancy for the drug-residue pair would be 150%. As seen in Figure 8b, this is what is happening in the highest occupancy case of CN045, forming hydrogen bonds with the sidechain of Asn382 (with an average occupancy of roughly 213%), and all other cases where occupancy exceeds 100%. At the same time, even in the cases where occupancy is below 100%, it could still be a sum of the individual donor-acceptor pairs present between the full molecular structures of the drug and residue pair. For clemastine, the highest occupancy is with Tyr106, at approximately 163% averaged across the replicas. It is worth mentioning that for many of these pairs, the inverted pair (where the donor and acceptors are flipped) is also present in the Top 20 occupancies. This may seem confusing, especially since it was established that both compounds only contain hydrogen bond acceptors. The reason for this is expected, since VMD by default considers any heavy atom covalently bound to a hydrogen as a donor if it also meets the cutoff criteria mentioned above. So, weakly electronegative donors such as carbon atoms are still represented in this analysis and there are instances where even the compounds appear as donors. For example, in the case of the M1-CN045 complex (Figure 8b, left), the top occupancy occurs when Asn382 is acting as the donor, but down the line there is a minor occupancy present when CN045 is acting as a donor instead. In this case, one of the aliphatic hydrogens on CN045 is likely interacting closely with the carbonyl oxygen on the sidechain of Asn382. Thus, the analysis is encompassing both the conventional polar hydrogen bonds as well as weaker interactions involving hydrogen.

Interestingly, there is a notable incidence of overlapping residues when comparing the occupancy plots of both M1 systems. Within the top 20, there are 7 dominating residues between both systems - both compounds maintain hydrogen bonds with Ala196, Asn382, Ser109, Trp157, Trp378, Tyr106, Tyr381. Four of these residues are also within the top 5 occupancies in both systems, namely Asn382, Ser109, Trp378, Tyr106. It is worth noting that nearly all of these residues were also the most pronounced cases where the RMSF was reduced when the compounds were bound to M1 (Figure 7c). In other words, these specific geometry-based hydrogen-bond-like contacts are consistent with, and may contribute to, the greater stability observed for the corresponding orthosteric-site residues.

With the H3-CN045 complex (Figure 8c, left), we see that hydrogen bond occupancies were significantly reduced, compared to the other two systems. The highest occupancy is observed with Arg381, at approximately 122% averaged across the replicas, although with a rather sizable standard error of nearly 55%. After this, all other occupancies drop significantly, beginning with hydrogen bonds with Tyr115, with an average occupancy of 65% +/- 38. It is worth noting that the hydrogen bond analysis is conducted over all residues interacting with the compounds (under the distance and geometric criteria mentioned previously), and not specifically with the pre-defined orthosteric residues of each system. This is why not all residues in this analysis are also included in the RMSF plots – the RMSFs were calculated specifically against an index group of known orthosteric residues, most of which were present and interacting with the compounds, in the initial docking configurations. As the simulations progress and the system evolves, the list of residues meeting the hydrogen bond criteria can naturally change over time. Given the instability in the H3-CN045 simulations, and the fact that in at least one replica, CN045 leaves the pocket entirely, there is much less overlap between the paired residues in this analysis and the formal orthosteric site residues. This is in contrast to the M1 complex simulations, where there is more consistency between the residue list. The pairs for which n=1 likely correspond to the pairs formed when CN045 was moving throughout the system and transiently interacting with residues well beyond the orthosteric site. At any rate, the residues here that are also in the RMSF analysis of the H3 orthosteric site (Figure 7d), have relatively low occupancies and also induced higher RMSFs. In other words, there were not enough persistent contacts consistently present between H3 and CN045, to stabilize the orthosteric site residues. Furthermore, when observing the moving average (across replicas) of the total number of hydrogen bonds for the H3-CN045 complex, this number falls to 10.45, much less than the figures for both M1 complexes.

### 3.6 Binding free energy analysis *ΔG_bind_* with MM-PBSA

#### 3.6.1 Comparing Total *ΔG_bind_* of all 3 systems

As a further measure of complex stability, end-state free-energy calculations were performed with MM-PBSA. This analysis provided a global view of binding energetics and also identified residues that contribute most strongly to stabilization or destabilization of the complexes.

Figure 9 summarizes the final *ΔG_bind_* values for each replica of each complex (left) together with the individual energy components contributing to those values (right). More negative *ΔG_bind_*values are generally interpreted as more favorable binding within the assumptions of MM-PBSA. Overall, the two M1 systems showed similar *ΔG_bind_*values across replicas: M1-Clemastine averaged -17.64 ± 2.86 kcal/mol, whereas M1-CN045 averaged -18.55 ± 1.96 kcal/mol. The H3-CN045 system was less favorable on average, at -14.39 ± 1.92 kcal/mol.

**Fig. 9:**
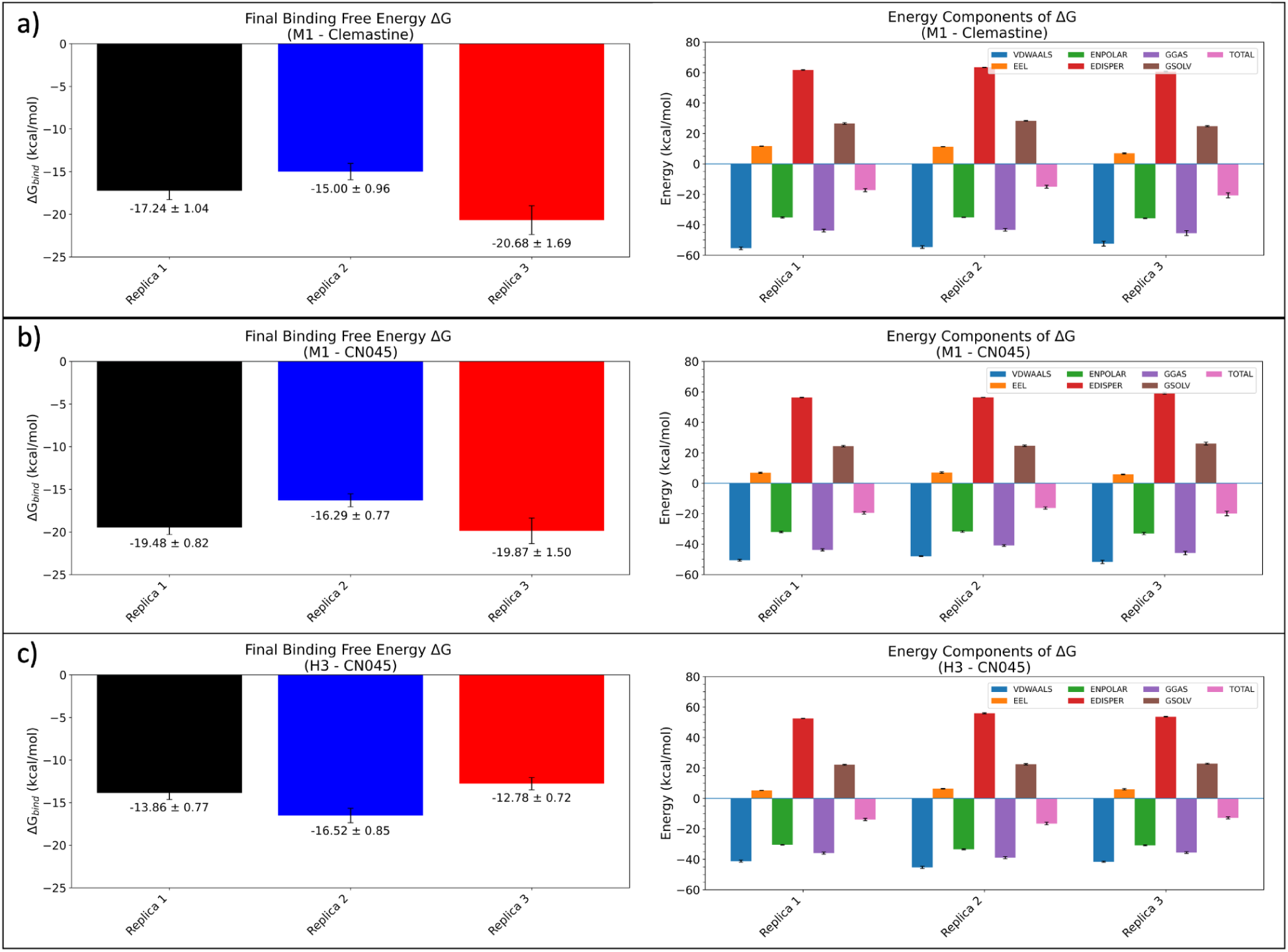
The final ΔG_bind_ for each replica of each system is illustrated on the left and the breakdown of energy components contributing to ΔG_bind_ on the right.

Both M1 systems share a similar pattern throughout the replicas, where the lowest energy is found with replica 3, followed by replica 1, and replica 2 having the highest energy. The pattern in the H3-CN045 complex was slightly inverted, with the lowest being replica 2, followed by replica 1, and replica 3 having the highest energy. These relative differences between respective replicas can be better understood by observing the difference in energy components (Figure 9, right). For example, replica 2 presenting the lowest energy in the H3-CN045 complex can be explained by having lower van der Waals energies (VDWAALS), gas phase interaction energy (GGAS), and non polar solvation energy (ENPOL). Another example is also replica 3 of the M1-Clemastine system, the lowest energy replica here. The component plot highlights a reduced gas phase interaction energy (GGAS), gas phase solvation energy (GSOLV), dispersion energy (EDISP), and electrostatic energy (EEL). Additionally, further information can be gathered by observing the per-residue decomposition plots in Figure S7, highlighting what residues contributed the most to a particular replica. Once again, in the case of replica 3 of M1-Clemastine, with a *ΔG_bind_* -20.68 +/- 1.69, we can see that this was driven largely by Tyr106 and Ser109, presumably from forming more favorable interactions with clemastine during these trajectories.

Nonetheless, at first glance, a few details can be observed in Figure 9. In the M1-CN045 system, all 3 replicas yield lower energies than ⅔ replicas of the M1-Clemastine system. Conversely, at least 1 replica in the latter has a lower energy than all 3 in the former. In both M1 systems across the board, however, energies are lower than all replicas in the H3-CN045 system. To further understand these variabilities, weighted and unweighted statistical analyses were conducted (Figure S8). In the unweighted analysis, M1-CN045 had a 55.6% probability of being a lower *ΔG_bind_* than M1-Clemastine, and an 88.9% probability of being lower than H3-CN045. These probabilities indicate only modest evidence for a difference with clemastine, and stronger evidence for a difference with H3-CN045.

In the weighted analysis, within-replica variability was incorporated through inverse-variance weighting, so replicas with smaller SD values contributed more heavily. Under this sensitivity analysis, the probability that M1-CN045 was lower than M1-Clemastine increased to approximately 94%, reflecting the smaller within-replica SDs for M1-CN045. The probability that M1-CN045 was lower than H3-CN045 approached 100% under the same assumptions. These results illustrate how incorporating within-replica variability can shift the inferred probabilities, especially when variability differs substantially between systems. Because these estimates are based on only three replicas per system and temporally correlated trajectories, they should be interpreted cautiously and primarily as comparative support for the broader conclusion that H3-CN045 is less stable than either M1 complex.

#### 3.6.2 Per-residue *ΔG_bind_* decomposition analysis of the M1 complexes

The analysis in the previous section provided a broad view of the thermodynamic stability of all three complexes and indicated that H3-CN045 was the least stable. Together with the other computational analyses, this finding helps explain why CN045 behaves as a weaker binder to H3 than to M1 in the binding assays. Given CN045’s preference for M1, and the likelihood that M1 is among the biologically relevant targets in the OPC-differentiation pathway, a more detailed free-energy analysis of the two M1 complexes was warranted. In this section, we examine per-residue contributions to total binding free energy, both overall and frame by frame, together with an energy-component breakdown.

Figure 10a illustrates the average contribution to *ΔG_bind_*from each residue, across all 3 replicas, for each M1 complex. They are sorted based on the descending order (right to left) in the M1-CN045 complex. The residues in the M1-Clemastine system are paired accordingly. This provides a more clear look at how these contributions differ between the systems. For clarity, data labels were only applied to residues which presented an energy contribution with an absolute value greater than or equal to 0.4 kcal/mol. A few things are clear from this analysis. For one, the most dominant stabilizing residues (negative energy) appear to be Trp378, Ala196, Tyr381, Tyr106, Gln110, Asn382, Val385, Trp157, and Ser109. In the vast majority of these residues, the energy was much more negative when bound to CN045, than when bound to clemastine. Within this subset of residues, in at least one case (Tyr381) there is a large positive energy (destabilizing) contribution when bound to clemastine. It is important to note that most of these stabilizing residues are also in the Top 20 hydrogen bond occupancy list (Figure 8), and were also some of the orthosteric residues that presented a reduced RMSF when bound to the drug (Figure 7). So, in other words, the increased structural stability (lower RMSF) in these complexes is consistent with the average measure of thermodynamic stability (*ΔG_bind_*) contributions per residue, and it can be traced back to the specific drug-residue interactions, such as some of the specific bonds that are driving much of this.

**Fig. 10:**
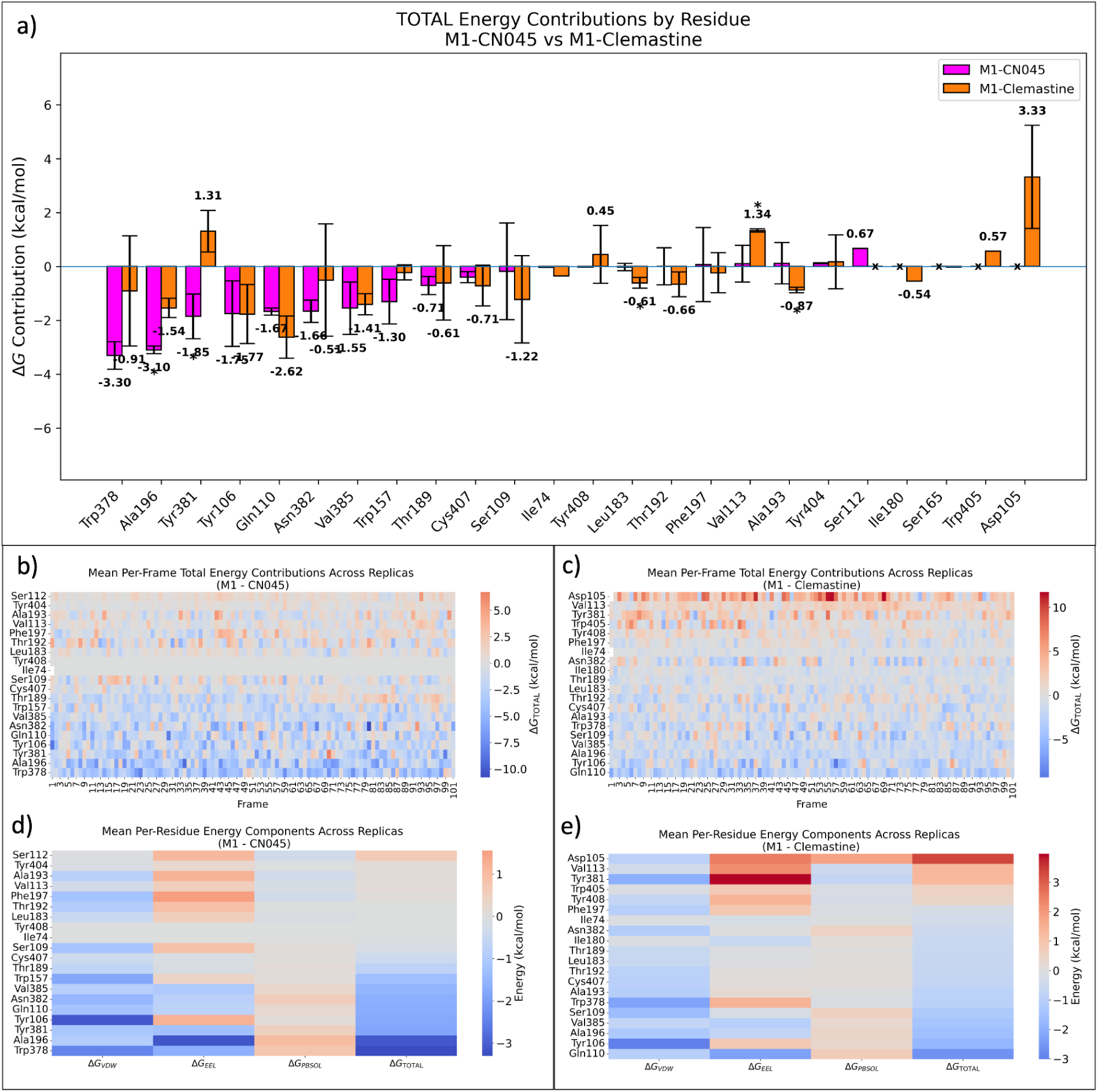
Per-residue ΔG_bind_ breakdown for the M1 systems. a) Total contributions per residue for each system. In this plot they are sorted based on the descending order of the M1-CN045 system and then paired accordingly. b) The average contribution to ΔG_bind_ from each residue, per frame. c) The average contributions per residue, to each energy component of ΔG_bind_, across all frames. All values are averaged across all 3 replicas in each system. Asterisks are placed on the most statistically significant pair differences (p-value <0.05, based on Welch’s t-test [77]), and X’s are placed for residues which did not contribute to ΔG_bind_ at all across the replicas of the respective system.

Still, the overall *ΔG_bind_* is more comparable between both systems than the difference presented in this subset of residues. This is, in part, because there are a few other residues that are specifically helping stabilize the M1-Clemastine complex, that are largely not involved in the M1-CN045 complex. These include Leu183, Thr192, Ala193, and Ile180. Another notable difference is that in the M1-Clemastine complex, there are many more destabilizing residues, making positive energy contributions. These include Tyr381, Tyr408, Val113, Trp405, and Asp105. On the other hand, there is only one destabilizing residue in the M1-CN045 complex with an energy contribution above 0.4 kcal/mol. That is Ser112, contributing an average of 0.67 kcal/mol. However, this only happened in one replica, which is why there is no error bar. It is also important to note the highly positive energy contributed by Asp105 in the M1-Clemastine complex. These aspartate residues are highly conserved across class A GPCRs, and play a crucial role in endogenous ligand binding (acetylcholine and histamine for the M1 and H3 receptors, respectively) and subsequent receptor activation. They include Asp105 in the M1 receptor [32,75] and Asp114 in the H3 receptor [33,76]. The fact that it produces such a positive energy contribution in at least ⅔ replicas (see Figure S7) indicates that clemastine may have a tendency to destabilize this residue. This is not the case with CN045, and given the highly important role of this residue, the simulations identify a potentially relevant difference between the compounds.

A statistical analysis was further performed employing Welch’s t-test [77], to help uncover the pairs of residues that are statistically different enough (p-value < 0.05), which takes into account variance between each system’s replicas and not just the mean values. These residue pairs are highlighted with an asterisk in Figure 10a. The analysis reveals that Ala196, Tyr381, Leu183, Val113, and Ala193 had the most statistically different contributions to each system’s *ΔG_bind_*. In ⅗ of these (Ala196, Tyr381, and Val113), the difference was a noticeably lower energy contribution when bound to CN045 over clemastine, while the opposite was noted for the remaining two residues (Leu183 and Ala193). It’s important to note that not all residues in Figure 10a contributed to each system’s *ΔG_bind_*. The bars in this plot represent data from at least one replica for each system. For the residues that did not contribute at all to the binding energy in any replica, an X was drawn where the bar plot would have been. For these pairs, *p*-values could not be calculated, however noticeable differences can still be observed (as seen with Asp105).

Figures 10b,c provide heat maps that show the residue decomposition in every frame of the MM-PBSA analysis. Once again, this corresponds to the last 100 frames of the 5000 frames (corresponding to the last 20 nanoseconds of the full 1 microsecond simulation) in the post-processed trajectories. They are averaged across replicas and sorted in descending order (top to bottom) in energy; the stabilizing residues are in the blue range while the destabilizing residues are in the light orange to red range. For the M1-CN045 complex, we observe the same set of stabilizing residues from Figure 10a (Trp378, Ala196, Tyr381, Tyr106, Gln110, Asn382, Val385, Trp157) in the bottom half of the heat map. Throughout most of the frames, these residues are contributing favorably to the complex (negative energy). There are some exceptions, however. For example, in frame 70, Tyr381 makes a highly positive contribution. The same can be observed in frames 75 and 92 of Asn382. This demonstrates the variation found throughout the frames and how the sampled configurations in the binding site can transiently shift the energy contributions. The same can be observed in the top half, where there are frames when these residues contribute favorably (blue), while the opposite being the case in the majority of the frames. The constant grey noticed with Ile74 and Tyr408 indicate that these residues almost negligibly contributed to the binding energy.

When observing the heat map for the M1-Clemastine system (Figure 10c), there is a clear difference. For one, there are many more frames where residues are making highly positive energy contributions. This is seen by the larger number of orange frames, as well as many frames colored in dark red. The bottom half also includes more orange frames, and less dark blue frames. Additionally, the contributions of Asp105 are rather clear, consistently contributing greater than +7.5 kcal/mol, and many instances where it is greater than +10 kcal/mol (frames 36, 55-57, and 70). The same trend can also be observed with Tyr381 and Val113, although with smaller magnitudes in energy. In essence, the data in these heat maps provides a frame-based representation of the bar plot in Figure 10a.

Lastly, the heat maps in Figure 10d,e provide a look at the per-residue contributions to the energy components of *ΔG_bind_*. There are similarities and differences here between the complexes. Both have predominantly negative van der Waals interactions (*ΔG_VDW_*), which is consistent with both compounds being largely hydrophobic and interacting with a largely hydrophobic binding site. In terms of electrostatic interactions (*ΔG_EEL_*), the majority of the residues yield positive contributions, implying the presence of repulsive charge-charge interactions. This appears to be more intensely captured in the M1-Clemastine complex, specifically with Asp105, Val113, and Tyr381. This suggests that the overall positive contributions of these residues, as observed in Figure 10a, are largely electrostatically driven – these residues and clemastine are intensely repelling through their partial atomic charges, with Tyr381 accounting for most of this. In both complexes, the polar solvation energy contributions (*ΔG_PBSOL_*) are largely neutral to positive, with Asp105 and Ala196 contributing the most in the clemastine and CN045 complexes, respectively. The last column of the heat map, (*ΔG_TOTAL_*) depicts the collective sum of these contributions per residue. In M1-CN045, the distribution transitions smoothly from a mildly positive contribution at Ser112 to strongly negative contributions from Tyr381, Ala196, and Trp378. In M1-Clemastine, the distribution is shifted toward more positive contributions, beginning with the strongly destabilizing Asp105 and followed by Val113 and Tyr381.

The decomposition analysis provides a more detailed thermodynamic picture of how each complex is stabilized. In particular, several destabilizing interactions are present in M1-Clemastine but not in M1-CN045, whereas several strongly favorable interactions are enhanced in M1-CN045. Although the global *ΔG_bind_*values of the two M1 complexes are relatively similar, the per-residue analysis clearly distinguishes them and suggests that the most consequential differences involve Asp105, Val113, Tyr381, Ala196, and Trp378.

### 3.7 Comparing simulation ensembles to active and inactive state M1 receptor conformations

Up to this point, analysis of the two M1 simulation sets (clemastine-bound and CN045-bound) has shown broadly similar ligand stability, slightly lower orthosteric-site RMSF for clemastine than for CN045, and a similar number of persistent strong and weak hydrogen-bond-like contacts (approximately 15). The binding free-energy calculations further indicate comparable overall *ΔG_bind_* values, although the per-residue decomposition analysis reveals potentially functionally relevant differences, particularly for Tyr381, Tyr404, and Asp105. A central question therefore remains: clemastine binds strongly to M1 (Figure S3b), yet CN045 is nearly twice as potent as clemastine in the OPC-differentiation assay. If M1 is a relevant target for both compounds, then differences in the receptor conformational ensembles stabilized by each ligand may help explain their divergent biological behavior. The working hypothesis in this manuscript is that inhibition of M1, rather than binding alone, contributes to OPC differentiation. In this section, we therefore examine simulation metrics that are commonly used as proxies for more active-like versus more inactive-like local conformations of M1, while recognizing that these structural signatures do not by themselves establish functional efficacy.

#### 3.7.1 Clemastine stabilizes an active-like NPxxY microswitch ensemble at Tyr^7.53^ (Tyr418)

The NPxxY motif is a highly conserved sequence (Asn–Pro–x–x–Tyr) located at the cytoplasmic end of TM7 (short for transmembrane helix 7) in class A GPCRs. The terminal tyrosine, Tyr418 (or Tyr^7.53^ in the Ballesteros–Weinstein numbering system) undergoes state-dependent rotameric rearrangements that are tightly coupled to receptor activation and intracellular signaling [78–80]. Across GPCR families, rearrangements at the NPxxY segment (specifically the Tyr418 residue in this case) accompany the transition toward active conformations and help stabilize the active intracellular architecture required for G protein binding. A commonly accepted method of measuring these conformational shifts is the χ1 rotamer dihedral angle of this residue. Comparisons can then be made against the rotamer’s known active-state and inactive-state dihedral angles. The reference angle for the active state was obtained from a recent cryo-EM structure (PDB: 9UCP) of the M1 receptor bound to a G protein and iperoxo (a known agonist), while the inactive state was measured from the original crystal structure used for these simulations (PDB: 5CXV [32]). Since χ1 rotamer populations are ensemble properties, the histogram view in Figure 11a is particularly informative: it reveals an ensemble-averaged representation of which basins are actually most populated, rather than relying on noisy frame-to-frame fluctuations. In the present simulations, clemastine exhibits a dominant Tyr418 χ1 rotamer basin that lies near the active-reference χ1 angle (Figure 11a), which is consistent with the replica-averaged time series showing sustained sampling closer to the active reference. In contrast, CN045 preferentially induces a distinct basin near ∼ −170° that is not aligned with either the active or inactive reference state, corresponding to a “non-productive” microswitch conformation, perhaps forming part of known non-productive metastable states observed in other GPCRs [81]. The next dominant angle induced by CN045 is around roughly 65°, in closer proximity to the inactive state reference conformation. While the clemastine simulations also present a peak at this angle, the probabilities are substantially lower.

**Fig. 11:**
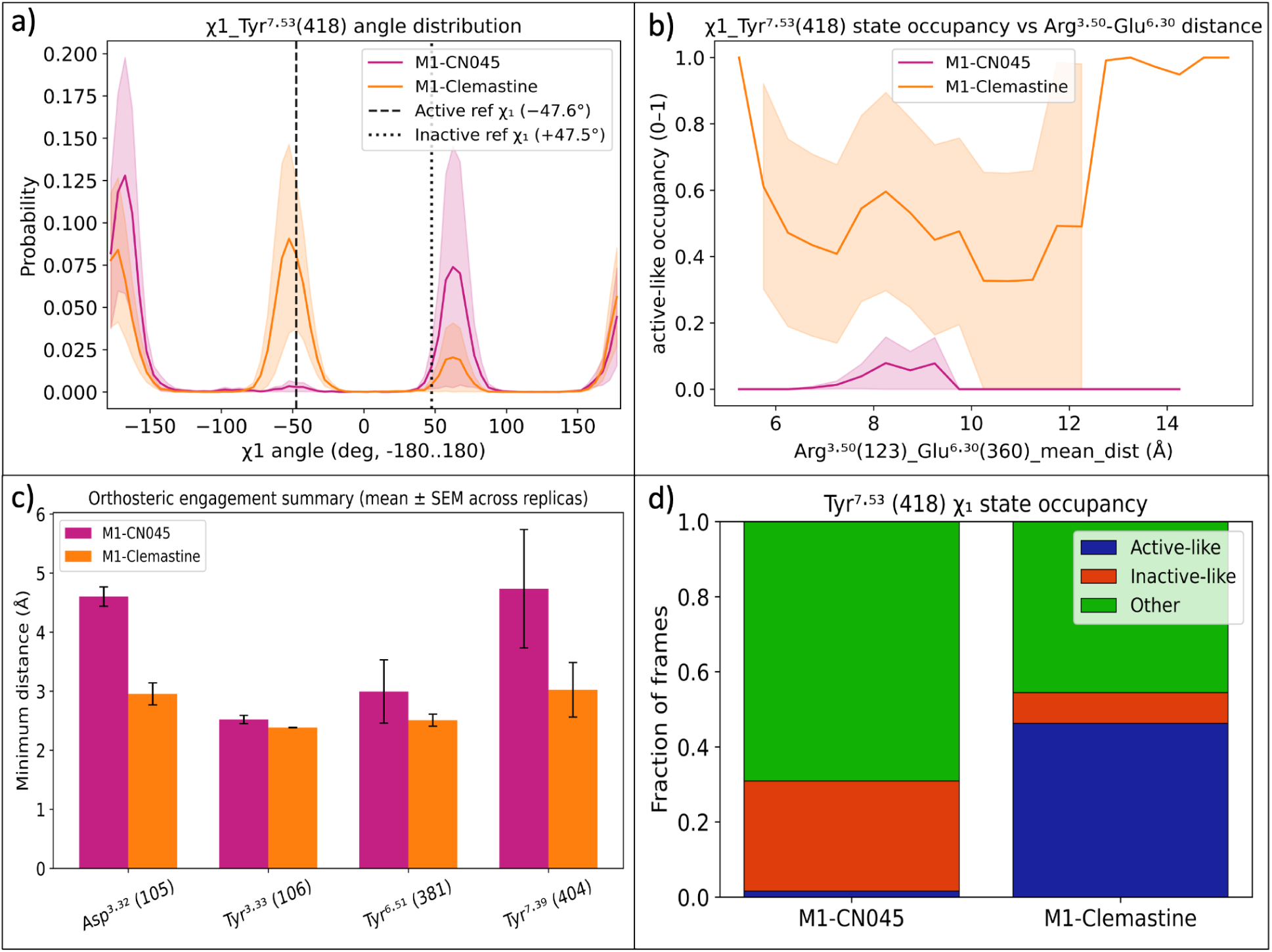
a) Histogram analysis of most dominant dihedral angles of χ1 observed in each system. b) Correlated map of active-like occupancy (based on being within a ± 25 degree threshold of the known active state χ1 of −47.6°) and binned distance of the “ionic lock” between Arg^3.50^(123) and Glu^6.30^(360). c) Minimum distances measured between each ligand and the key orthosteric residues (Asp105 and the aromatic cage residues Tyr106, Tyr381 and Tyr404). d) Overall summary of state occupancy as a fraction of frames in the two M1 systems.

A more global look at this data is presented in Figure 11d, where the occupied rotamer states of Tyr418 are shown with respect to the fraction of the frames occupying them. This plot highlights the clear divide in the rotamer populations between both systems. When bound to CN045, Tyr418 adopts a conformation that is neither considered active nor inactive, approximately 69.1% of the time (green). On the other hand, approximately 29.3% (red) of the frames correspond to the inactive state, and a mere 1.6% to the active state. Conversely, a very different distribution is observed when bound to clemastine. In this case, 46.3% of the frames demonstrate Tyr418 occupying the active state, while 8.2% of the frames are spent in the inactive state, and roughly 45.5% in neither. In terms of this specific rotamer conformation and its known role as part of a “microswitch” between states, the data indicates that CN045 more often samples inactive-like or non-productive Tyr418 χ1 states in this analysis, while the opposite is seen with clemastine.

It is important to note that this is not asserting that the receptor becomes fully active under the influence of clemastine in these trajectories, especially since the simulations were built from the 5CXV structure to begin with (which itself is in the inactive state) and the time scales to reach full activation would likely exceed 1 microsecond. Rather, the χ1 distributions indicate a relative bias: clemastine shifts the NPxxY/Tyr^7.53^ microswitch ensemble toward an active-like basin, when compared to CN045. Over time, inducing this active-like conformation at Tyr418 could help lower the energy barriers associated with the inactive-to-active state transitions. Such ligand-dependent shifts in conserved microswitch ensembles are consistent with the modern view of GPCR efficacy as stabilization of distinct conformational ensembles rather than a binary inactive/active switch [78].

#### 3.7.2 Clemastine couples NPxxY behavior to weakening of the DRY/TM6 ionic-lock contact

Another independent activation proxy comes from the intracellular DRY motif on TM3 (transmembrane helix 3) and its interaction network with TM6 (transmembrane helix 6). The DRY motif is a three-residue sequence (Aspartate(D)-Arginine(R)-Tyrosine(Y)) at positions 3.49, 3.50, and 3.51, respectively. Many class A GPCRs exhibit an inactive-state salt-bridge network involving Arg^3.50^ (Arg123 in the DRY motif of TM3) and Glu^6.30^ (Glu360 on TM6), often termed the ‘ionic lock’ (see Figure 12) [82]. Disruption of this interaction and other DRY motif rearrangements is commonly associated with TM6 outward movement and formation of the active intracellular binding site [83–86]. Here, Arg^3.50^–Glu^6.30^ distance serves as a direct, system-specific monitor of this interaction. Clemastine induced a larger mean separation than CN045, consistent with reduced ionic-lock engagement (see Figure 12 as an example observed in replica 1 of these simulations).

**Fig. 12:**
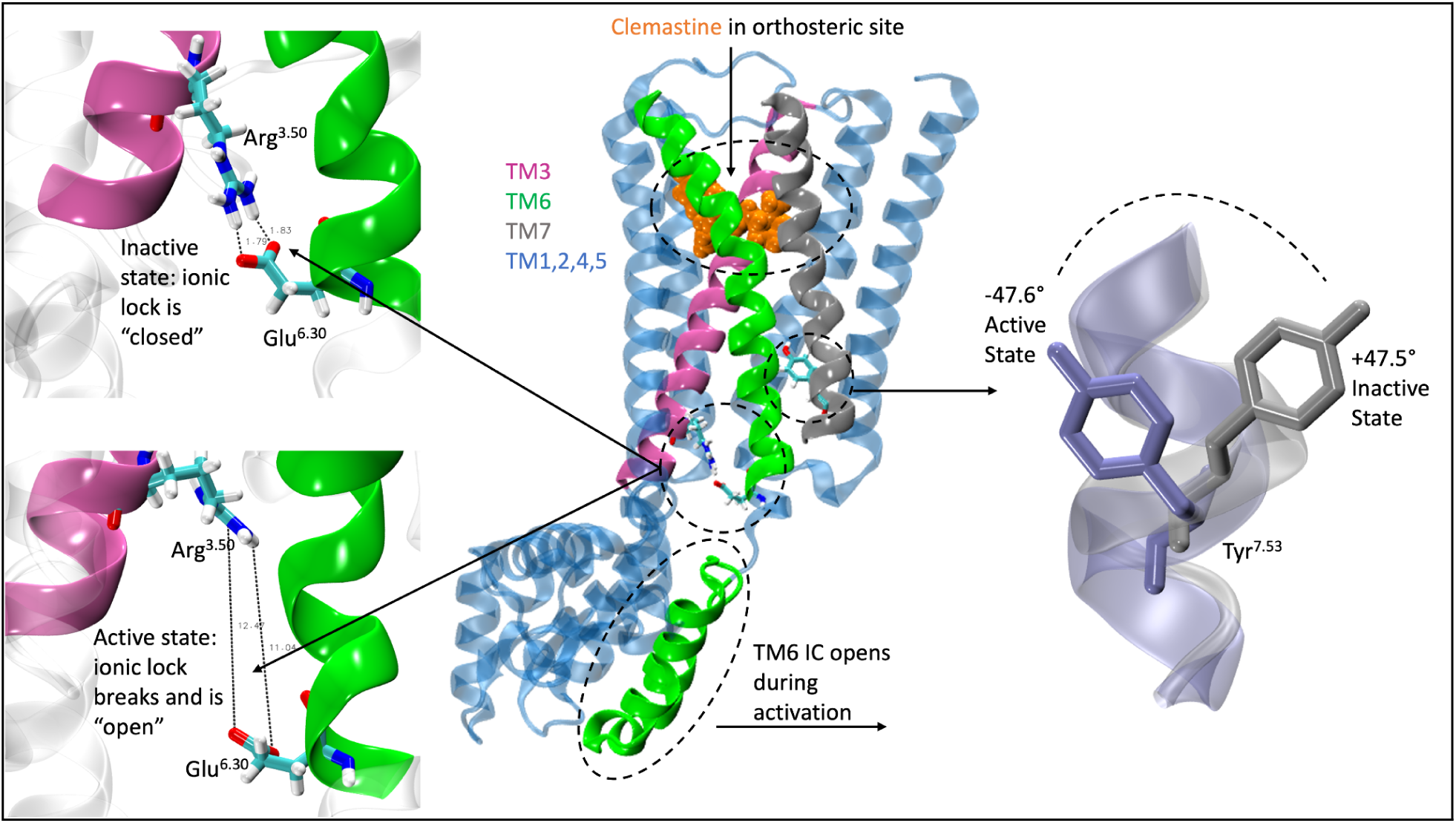
A closer look at the microswitch network. Clemastine binding is associated with a larger ionic-lock distance, an active-like conformational feature that is often associated with outward movement of the intracellular end of TM6, a precursor to G-protein engagement. Although the intracellular region in the 5CXV crystal structure contains a T4 lysozyme/endolysin fusion used for crystallization, the region highlighted here approximates the location of the TM6 intracellular end in the full-length receptor. Another hallmark of the active state is shown on the right: the change in Tyr7.53 χ1 from +47.5° in the inactive state to -47.6° in the active state.

More importantly, however, clemastine also shows mechanistic coupling between this ionic-lock proxy and the NPxxY microswitch mentioned above: an active-like Tyr^7.53^ χ1 occupancy increases at larger Arg^3.50^–Glu^6.30^ distances (Figure 11b). Here, the lines correspond to the mean value across each system’s replica simulations, and the shaded bands capture the variance across replicas. Regions with no shading indicate only one replica demonstrates this coupling behavior in that range. This binned relationship supports a coherent activation pathway interpretation: when the DRY/TM6 ionic contact loosens, consistent with one feature often associated with intracellular opening [82,87,88], the NPxxY/Y7.53 microswitch is more frequently in the active-like basin. CN045, in contrast, shows minimal active-like χ1 occupancy across distance bins, indicating weak coupling to this well-known activation-associated intracellular rearrangement. The opposite behavior can be seen in Figure S11, which shows this coupling behavior with respect to the *inactive* state conformation of the NPxxY microswitch. Here, an inactive-like Tyr^7.53^ χ1 occupancy increases more so for CN045 at larger Arg^3.50^–Glu^6.30^ distances. Together, this data indicates that there is stronger mechanistic coupling towards the active-like local signatures for the M1-Clemastine ensembles, and towards the inactive-like for M1-CN045.

#### 3.7.3 Clemastine exhibits tighter orthosteric engagement at conserved pocket residues

Although global binding free energies and orthosteric RMSF may be similar between CN045 and clemastine, orthosteric geometry can still differ in functionally meaningful ways. Muscarinic receptors share a conserved orthosteric interaction network in which a cationic ligand moiety commonly forms an ion pair with Asp^3.32^ (Asp105), and the ligand is often cradled within an aromatic environment that includes Tyr^3.33^ (Tyr106), Tyr^6.51^ (Tyr381), and Tyr^7.39^ (Tyr404). Structural analyses of muscarinic receptor complexes describe this Asp^3.32^ ion pair and aromatic cage as key determinants of orthosteric recognition [32,89,90]. In the present simulations, clemastine maintains shorter minimum distances to these four residues than CN045 does (Figure 11c), indicating a more engaged orthosteric pose. In particular, the difference is much more pronounced for Asp105 and Tyr404. This increased engagement is also consistent with the overall and per-residue RMSF (seen in Figure 7) being lower for the orthosteric site when clemastine is bound. Interestingly, in 3 out of 4 of these residues, the free energy contributions are noticeably different (Figure 10a), with largely positive energy contributions observed when bound to clemastine.

These orthosteric differences provide a plausible upstream origin for the divergent microswitch and ionic-lock signatures. In other words, even subtle shifts in how the ligand packs against conserved orthosteric anchors can bias the receptor’s internal activation network, leading to different conformational ensembles. This is consistent with broad GPCR structure–function literature emphasizing that ligand efficacy often reflects how a ligand stabilizes the receptor’s conformational landscape (ensemble bias), rather than simply overall affinity.

#### 3.7.4 Overall interpretation

Taken together, the combined evidence supports a consistent directionality: clemastine samples more active-like local M1 microswitch configurations than CN045 in these simulations in that it (i) stabilizes an active-like Tyr^7.53^/NPxxY microswitch χ₁ basin, (ii) preferentially samples larger DRY/TM6 separations consistent with reduced ionic-lock engagement, and (iii) maintains tighter packing against conserved orthosteric pocket residues that participate in productive ligand recognition and can facilitate downstream coupling to activation-associated microswitches. This mechanistic differentiation is particularly valuable in the context of otherwise similar simulation observables (binding free energies, RMSF, etc.), highlighting that the key functional differences likely reside in how each ligand biases conserved activation pathways rather than in gross stability or affinity metrics.

## 4 Discussion and Conclusion

Demyelination remains a major pathological feature of several neurological disorders, including multiple sclerosis. Compounds that promote OPC differentiation therefore represent an attractive route toward next-generation remyelination therapies. Using molecular dynamics simulations, we investigated structural hypotheses regarding the mechanisms of action of two such compounds: the well-studied clemastine and our current lead compound, the publicly available CN045.

While clemastine was already known to engage the M1 receptor, much of our work focused on identifying plausible targets for CN045. Initial screening implicated both M1 and H3, with follow-up binding assays showing a markedly stronger preference for M1. The computational analysis presented here provides a structural rationale for those binding data. In particular, comparative simulations of CN045 bound to M1 and H3 showed that the ligand was substantially less stable in the H3 complex. In one H3 replica, CN045 left the binding pocket entirely, which was not observed in the M1 simulations. Orthosteric-site RMSF was also higher in H3 than in M1, and the number of persistent hydrogen-bond-like contacts was reduced. Finally, MM-PBSA estimates were less favorable for H3 than for M1, with the unweighted resampling analysis assigning an approximately 89% probability that M1-CN045 has a lower *ΔG_bind_* than H3-CN045. Taken together, these analyses support the interpretation that CN045 forms a less stable complex with H3 than with M1, consistent with the binding assays.

The second objective was to compare the M1-CN045 and M1-Clemastine complexes in an effort to identify differences that may contribute to their divergent biological effects on OPC differentiation. Overall, the two complexes were more similar to each other than either was to H3-CN045. The ligand-centered structural analyses indicated comparably stable trajectories, and the RMSF analysis showed reduced orthosteric-site fluctuation in both cases, with slightly lower RMSF for clemastine. The hydrogen-bonding analysis also showed that both systems maintained a similar number of contacts overall (15.58 versus 15.64), with seven residues-Ala196, Asn382, Ser109, Trp157, Trp378, Tyr106, and Tyr381-consistently contributing to ligand stabilization. Global *ΔG_bind_* values were likewise similar, although the unweighted analysis gave M1-CN045 a modest advantage. The per-residue decomposition, however, revealed a clearer distinction: many of the same residues contributed favorably in both systems, but those contributions were often more favorable for CN045, whereas clemastine showed several unfavorable contributions, including at Asp105. When combined with the microswitch analyses below, these data are more consistent with CN045 sampling less active-like-or relatively more inactive-like-local M1 ensembles than clemastine.

Further crucial details were uncovered when comparing the simulation ensembles to the active and inactive state structures of the M1 receptor. The Tyr^7.53^ χ1 rotamer, ionic-lock, and orthosteric packing analyses provide convergent, mechanistically interpretable differences between both drugs. Clemastine preferentially stabilizes an active-like NPxxY/Tyr^7.53^ microswitch basin and exhibits stronger coupling to DRY/TM6 disengagement, while simultaneously showing tighter engagement with conserved orthosteric anchors (Asp^3.32^ and the Tyr^3.33^/Tyr^6.51^/Tyr^7.39^ aromatic cage). These results support the conclusion that clemastine more strongly biases the receptor toward active-like conformational ensembles than CN045, offering a plausible structural rationale for their divergent biological effects.

The available experimental data show that both compounds promote OPC differentiation, and the combined computational and binding results presented here provide a plausible structural framework for interpreting that activity. The overlap in key molecular features and broad pharmacokinetic properties between CN045 and clemastine is consistent with some shared target engagement. At the same time, the simulations highlight important differences that may influence not only affinity for M1 but also the conformational ensembles sampled by the receptor, with clemastine favoring more active-like local signatures than CN045. This pattern is consistent with the working hypothesis that CN045 and clemastine bias M1 differently, but does not establish relative functional inhibition in the absence of direct signaling assays. These molecular features may help inform the development of a future pharmacophore for remyelination-promoting compounds. Additional experimental work, particularly direct functional assays that quantify antagonism, inverse agonism, or pathway bias, will be important to test these mechanistic hypotheses.

## Data Availability

All data, analysis, Galaxy workflows, and associated Python scripts for analyzing and plotting the data can be found at this GitHub repository https://github.com/BlankenbergLab/OPCdiff-TargetAnalytics.

## Author Contributions

BR developed and executed the computational framework and simulations for this project and wrote the manuscript. FC co-developed the Python scripts used in the final data analysis. JJ assisted in developing the analyses used to distinguish active-like from inactive-like states. WM helped set up the initial binding free-energy calculations and optimized the GROMACS simulations for faster runtime. SM oversaw the binding assays and initial target-identification studies. YY conducted the in vitro OPC-differentiation experiments for both compounds. BT and DB served as principal investigators and oversaw all aspects of the work. All authors read and approved the final version of the manuscript.

## Supplementary Information

**Table S1:** Full analysis of physicochemical and ADME properties for (a) clemastine and (b) CN045 as conducted with SwissADME [74].

|  | Clemastine | CN045 |
| --- | --- | --- |
| **Physicochemical Properties** |  |  |
| Formula | C21H26ClNO | C16H22Cl3NO2 |
| Molecular weight (g/mol) | 343.89 | 366.71 |
| # of heavy atoms | 24 | 22 |
| # of aromatic heavy atoms | 12 | 6 |
| Fraction of sp3 carbons | 0.43 | 0.62 |
| # of rotatable bonds | 6 | 7 |
| # of H-bond acceptors | 2 | 3 |
| # of H-bond donors | 0 | 0 |
| Molar refractivity | 105.03 | 97.01 |
| TPSA (Å<sup>2</sup>) | 12.47 | 21.7 |
| **Lipophilicity** |  |  |
| iLOGP | 4 | 4.37 |
| XLOGP3 | 5.03 | 4.95 |
| WLOGP | 4.62 | 4.39 |
| MLOGP | 4.18 | 3.69 |
| SILICOS-IT | 5.15 | 5.03 |
| Consensus Log P<sub>o/w</sub> | 4.59 | 4.49 |
| **Water Solubility** |  |  |
| Log S (ESOL) | -5.12 | -4.97 |
| Solubility (mg/ml; mol/l) | 2.64e-03; 7.67e-06 | 3.91e-03; 1.07e-05 |
| Class | Moderately soluble | Moderately soluble |
| Log S (Ali) | -5.03 | -5.14 |
| Solubility (mg/ml; mol/l) | 3.19e-03; 9.27e-06 | 2.64e-03; 6.68e-07 |
| Class | Moderately soluble | Moderately soluble |
| Log S (SILICOS-IT) | -7.08 | -6.17 |
| Solubility (mg/ml; mol/l) | 2.85e-03; 8.28e-08 | 2.45e-04; 6.68e-07 |
| Class | Poorly soluble | Poorly soluble |
| **Pharmacokinetics** |  |  |
| GI absorption | High | High |
| BBB permeant | Yes | Yes |
| P-gp substrate | No | No |
| CYP1A2 inhibitor | No | Yes |
| CYP2C19 inhibitor | No | No |
| CYP2C29 inhibitor | No | No |
| CYP2D6 inhibitor | Yes | Yes |
| CYP3A4 inhibitor | Yes | No |
| Log K<sub>p</sub> (skin permeation; cm/s) | -4.83 | -5.02 |
| **Druglikeness** |  |  |
| Lipinski | Yes; 1 violation: MLOGP>4.15 | Yes; 0 violations |
| Ghose | Yes | Yes |
| Veber | Yes | Yes |
| Egan | Yes | Yes |
| Muegge | No; 1 violation: XLOGP3>5 | Yes |
| Bioavailability Score | 0.55 | 0.55 |
| **Medicinal Chemistry** |  |  |
| PAINS | 0 alert | 0 alert |
| Brenk | 0 alert | 0 alert |
| Leadlikeness | No; 1 violation: XLOGP3>3.5 | No; 2 violations: MW>350; XLOGP3>3.5 |
| Synthetic accessibility | 3.41 | 2.67 |

**Table S2:** Summary table of ADME data highlighting adherence to Lipinski’s Rule of 5 parameters.

| Molecule | Formula | MW (g/mol) | #Heavy atoms | #Rotatable bonds | #H-bond A | #H-bond D | Log P | Log S | GI absorption |
| --- | --- | --- | --- | --- | --- | --- | --- | --- | --- |
| CN045 | C<sub>16</sub>H<sub>22</sub>Cl<sub>3</sub>NO<sub>2</sub> | 366.71 | 22 | 7 | 3 | 0 | 4.49 | -4.97 | High |
| Clemastine | C<sub>21</sub>H<sub>26</sub>ClNO | 343.89 | 24 | 6 | 2 | 0 | 4.59 | -5.12 | High |

**Fig. S1:**
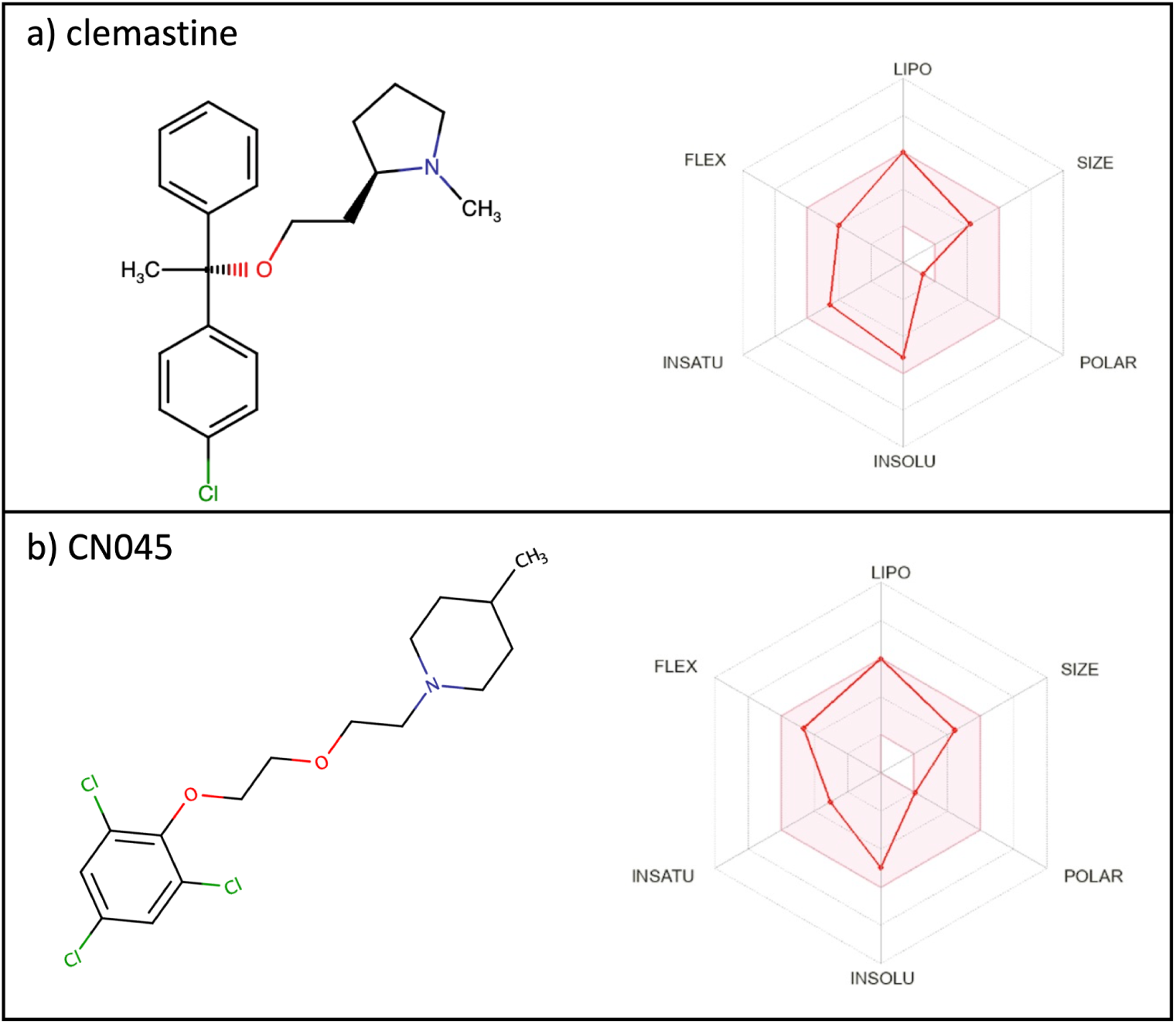
Molecular structures of both compounds along with a 3D distribution of ADME properties.

**Fig. S2:**
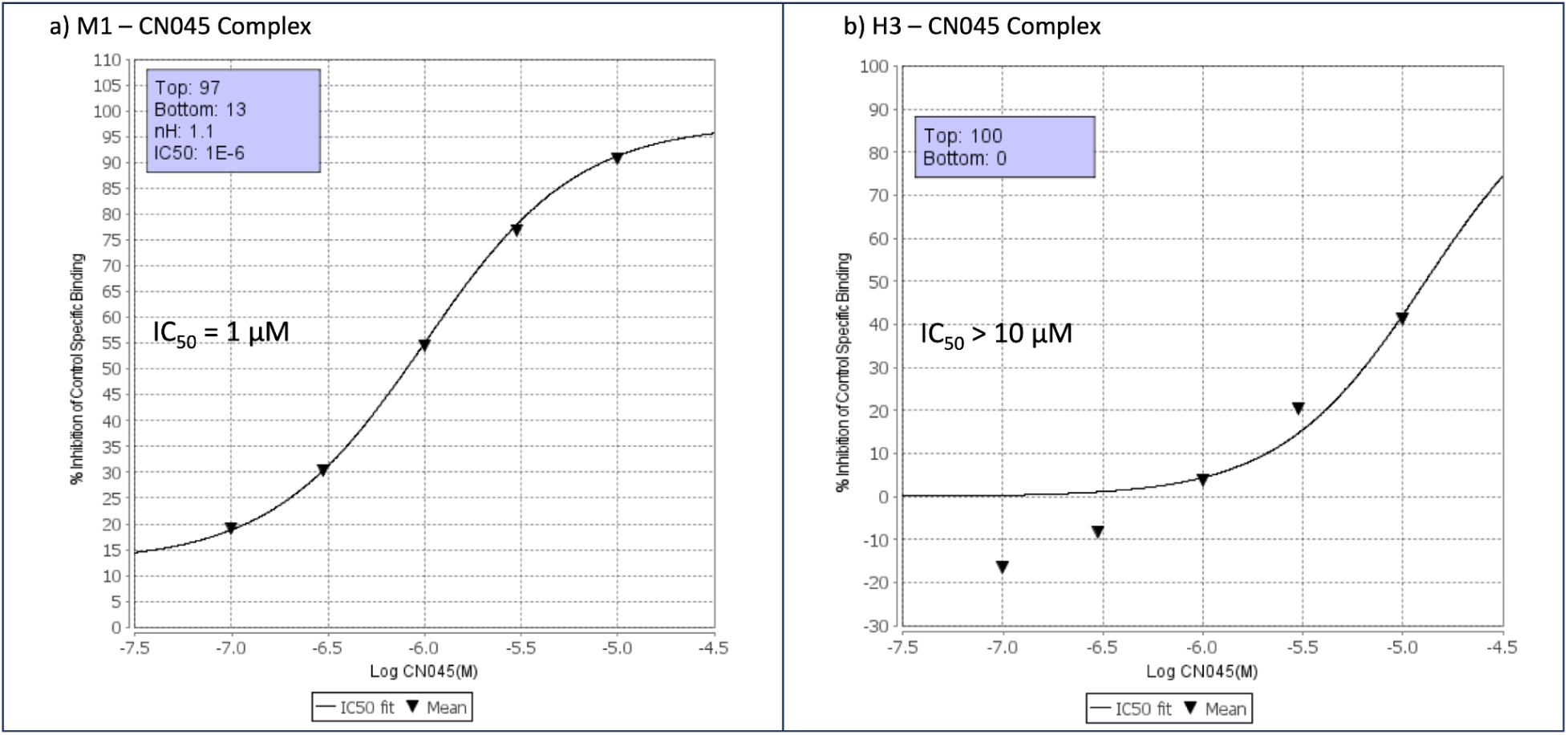
Binding assay plots of CN045 with the a) M1 muscarinic receptor and b) H3 histamine receptor.

**Table S3:**
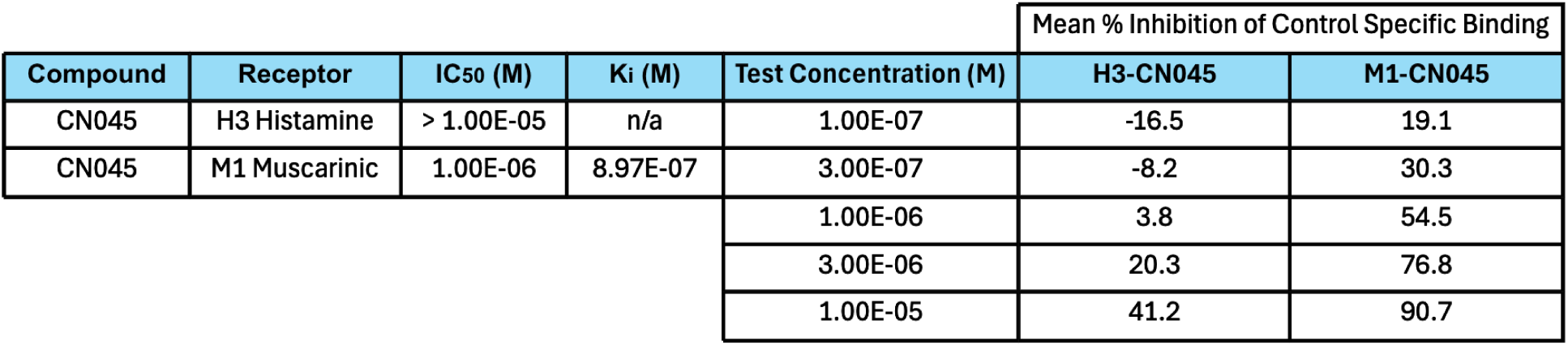
Summary of binding assay data for CN045 with the M1 muscarinic receptor and H3 histamine receptor. Mean % inhibition is measured at different tested concentrations, with the IC_50_ corresponding to the concentration where % inhibition surpasses 50%. In each experiment and if applicable, the reference compound (pirenzepine for M1 and (R)α-Me-histamine for H3) was tested concurrently with CN045. Compound binding was calculated as a % inhibition of the binding of the radiolabeled reference compound known to specifically bind to each respective receptor.

|  |  |  |  |  | Mean % Inhibition of Control Specific Binding |  |
| --- | --- | --- | --- | --- | --- | --- |
| Compound | Receptor | IC<sub>50</sub> (M) | K<sub>i</sub> (M) | Test Concentration (M) | H3-CN045 | M1-CN045 |
| CN045 | H3 Histamine | > 1.00E-05 | n/a | 1.00E-07 | -16.5 | 19.1 |
| CN045 | M1 Muscarinic | 1.00E-06 | 8.97E-07 | 3.00E-07 | -8.2 | 30.3 |
|  |  |  |  | 1.00E-06 | 3.8 | 54.5 |
|  |  |  |  | 3.00E-06 | 20.3 | 76.8 |
|  |  |  |  | 1.00E-05 | 41.2 | 90.7 |

**Fig. S3:**
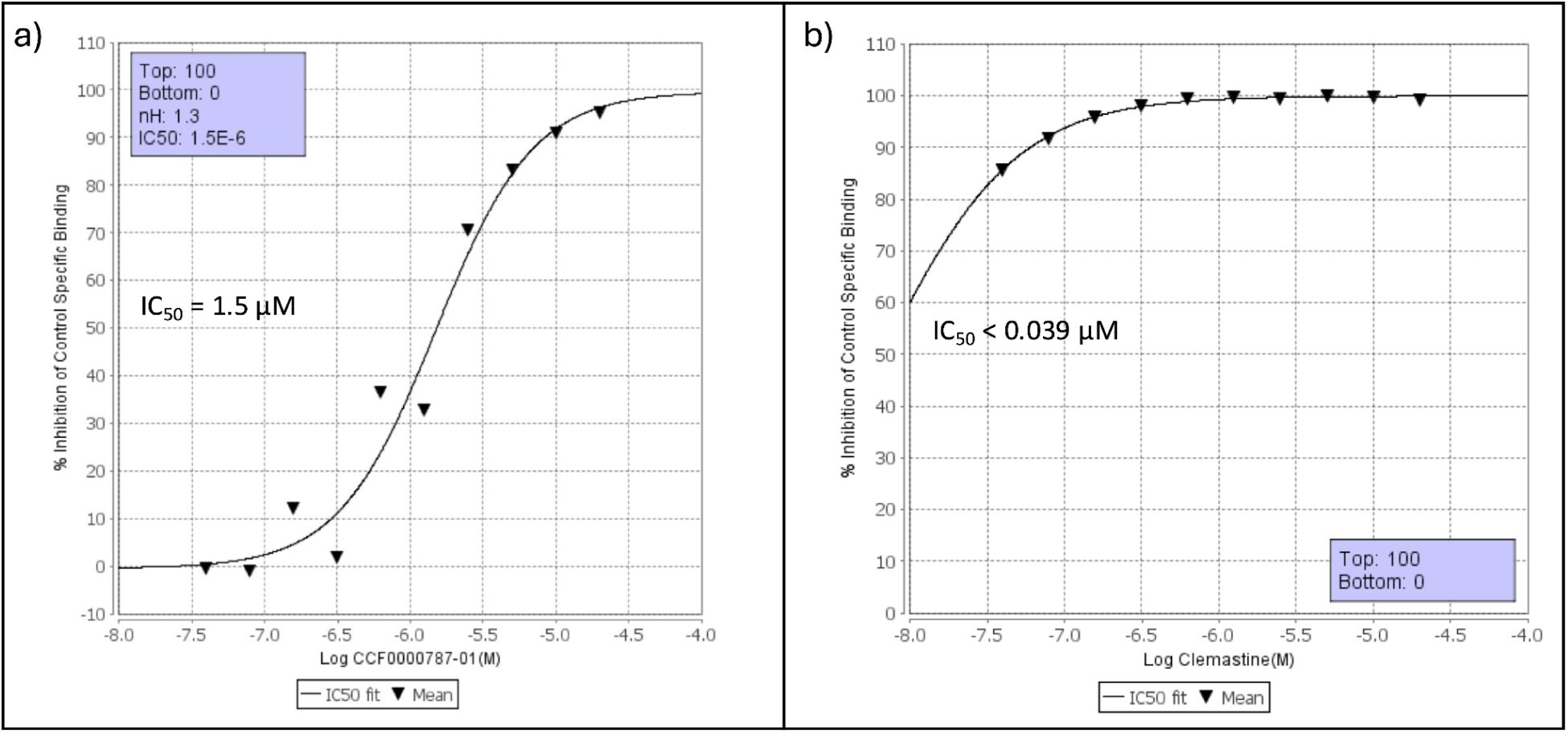
Binding assay plots of (a) CN045 (referred to as CCF0000787-01 in the plot) and (b) clemastine with the M1 muscarinic receptor.

**Table S4:** Summary of binding assay data for CN045 and clemastine with the M1 muscarinic receptor. Mean % inhibition is measured at different tested concentrations, with the IC50 corresponding to the concentration at which % inhibition exceeds 50%. In each experiment, where applicable, the reference compound (pirenzepine) was tested concurrently with CN045 and clemastine. Compound binding was calculated as the % inhibition of binding of a radiolabeled ligand known to bind specifically to the M1 muscarinic receptor.

|  |  |  |  |  | Mean % Inhibition of Control Specific Binding |  |
| --- | --- | --- | --- | --- | --- | --- |
| Compound | Receptor | $IC_{50}$ (M) | $K_i$ (M) | Test Concentration (M) | M1-CN045 | M1-Clemastine |
| CN045 | M1 Muscarinic | 1.50E-06 | 8.97E-07 | 3.91E-08 | -0.4 | 85.7 |
| Clemastine | M1 Muscarinic | < 3.9E-08 | n/a | 7.81E-08 | -0.9 | 91.8 |
|  |  |  |  | 1.56E-07 | 12.2 | 95.8 |
|  |  |  |  | 3.13E-07 | 2 | 98.1 |
|  |  |  |  | 6.25E-07 | 36.6 | 99.4 |
|  |  |  |  | 1.25E-06 | 32.8 | 99.8 |
|  |  |  |  | 2.50E-06 | 70.5 | 99.5 |
|  |  |  |  | 5.00E-06 | 83.3 | 100 |
|  |  |  |  | 1.00E-05 | 91 | 99.9 |
|  |  |  |  | 2.00E-05 | 95.4 | 99.3 |

**Figure S4:**
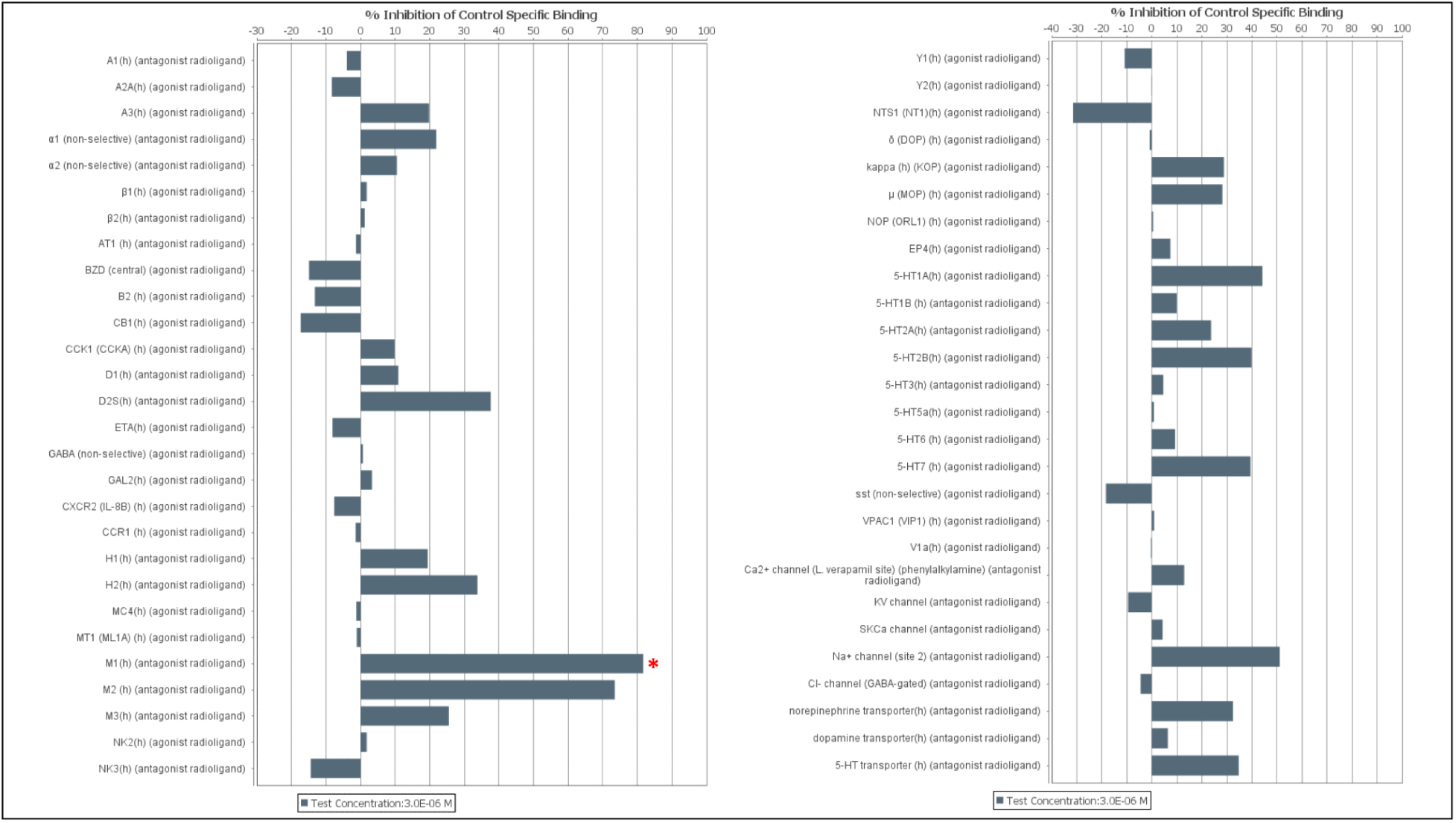
Results from the CEREP Express Panel conducted by Eurofins. These experiments suggested that CN045 produced the strongest inhibition of the M1 muscarinic receptor among the 55 targets screened and therefore helped prioritize M1 as a candidate target in the OPC-differentiation pathway.

**Figure S5:**
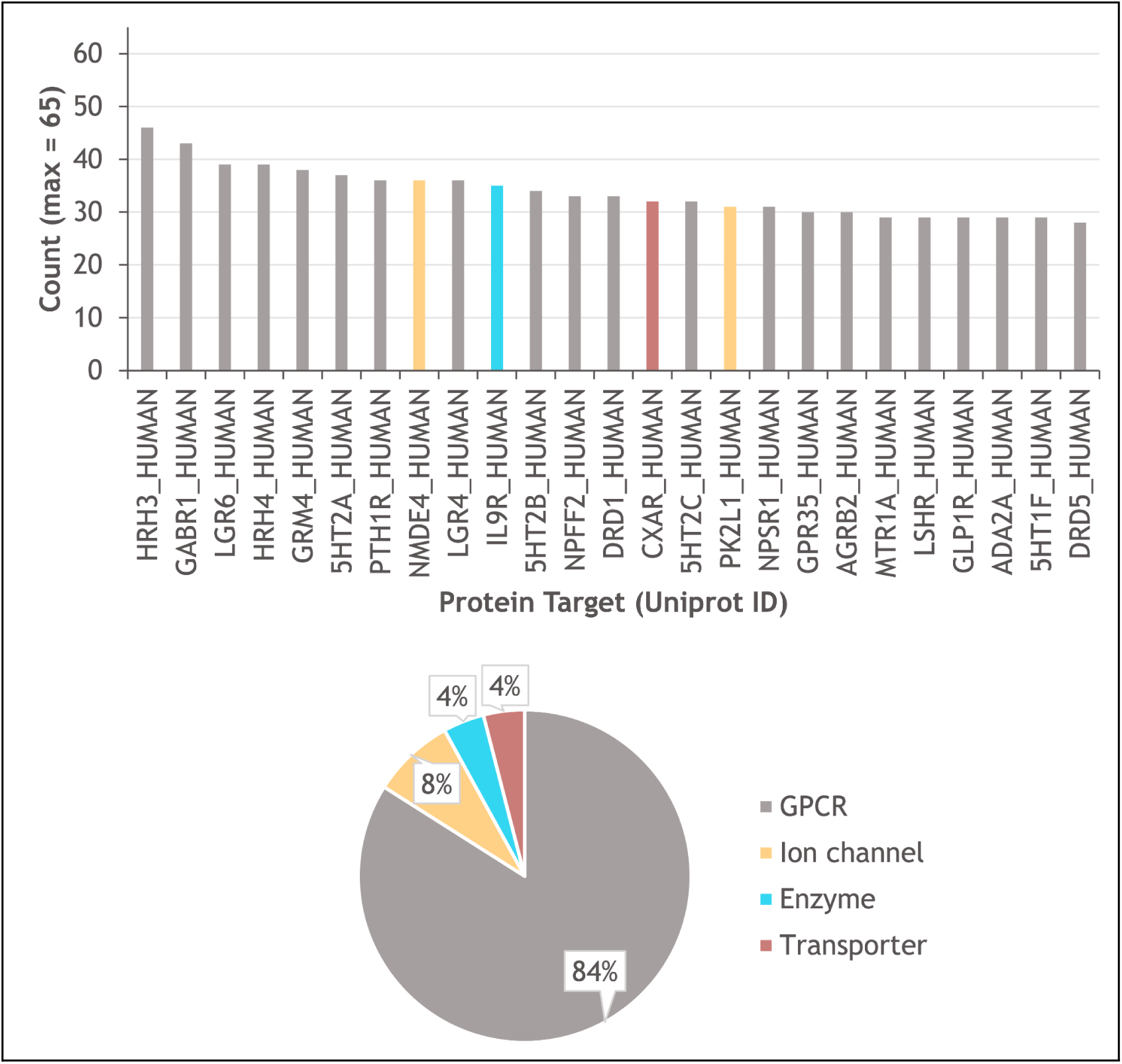
Results from a cheminformatics analysis conducted by Bristol Myers Squibb, using the Matchmaker software.

**Fig. S6:**
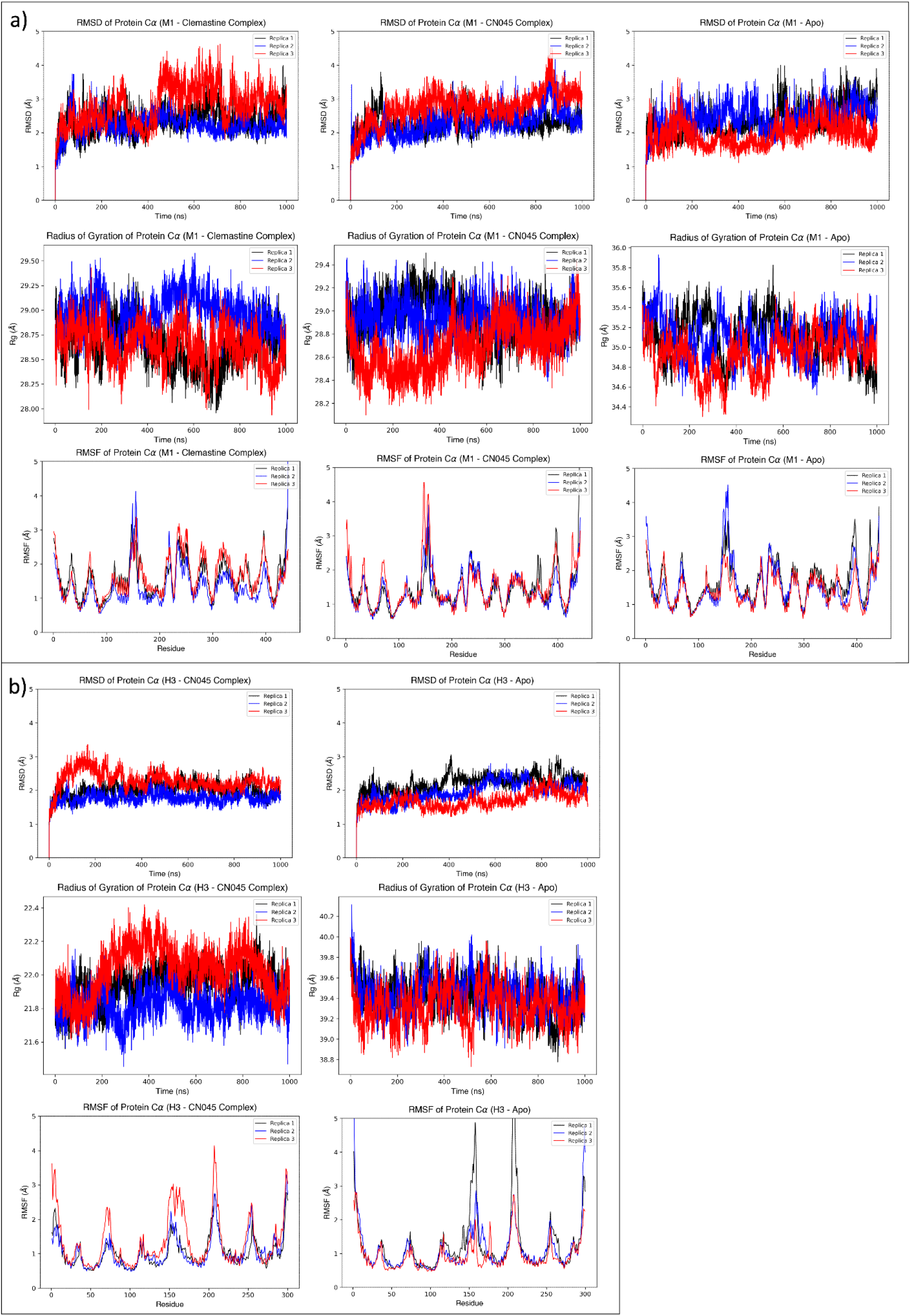
Structural analysis of overall protein structure as measured with RMSD, RMSF, and R_g_.

**Fig. S7:**
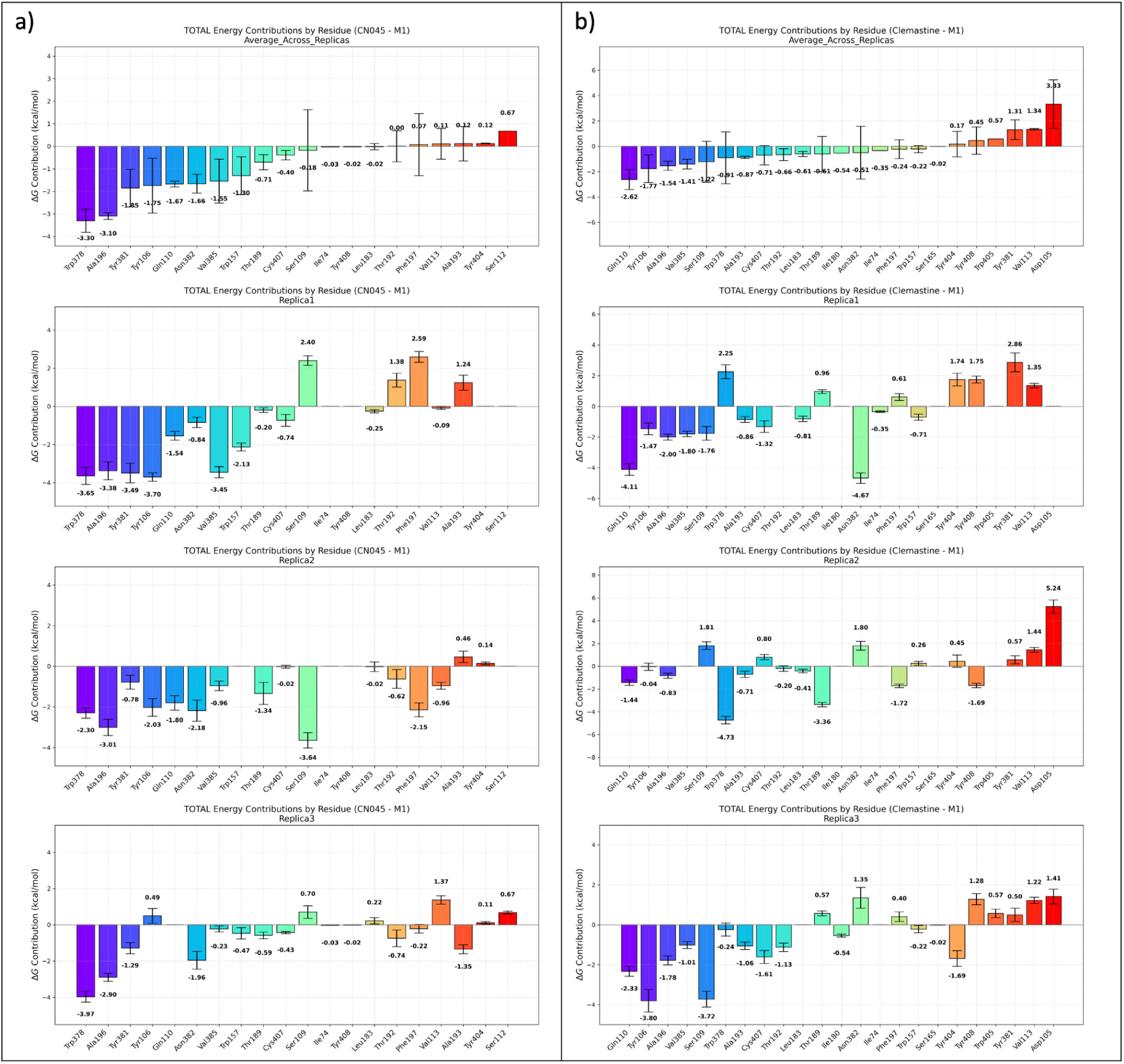
Decomposition of ΔG_bind_ per residue. The amounts represented here represent TOTAL energy contributions; for each residue, the energy is the energy difference between the complex, and the sum of the energies in the receptor and ligand. The final, across replica averages are sorted in descending order from right to left for each system, and are included at the top row. The energy decomposition of each replica is included in the bottom 3 rows. For each system, the replicas are sorted in the same order as the order in the top plot. This allows for a more clear comparison of how the individual replica decompositions vary relative to their aggregated, sorted order.

**Table S5:** Weighted and unweighted probabilities of M1-Clemastine and H3-CN045 yielding a lower binding free energy than M1-CN045.

| System | Unweighted Mean $\Delta G$ | Unweighted SD | Weighted Mean $\Delta G$ | Weighted SE (of mean) | Unweighted probabilities of lower $\Delta G$ | Weighted probabilities of lower $\Delta G$ |
| --- | --- | --- | --- | --- | --- | --- |
| M1-Clemastine | -17.64 | 2.86 | -16.72 | 0.65 | 44.40% | 5.80% |
| M1-CN045 | -18.55 | 1.96 | -18.04 | 0.53 |  |  |
| H3-CN045 | -14.39 | 1.92 | -14.18 | 0.45 | 11.10% | 0.00% |

**Figure S8:**
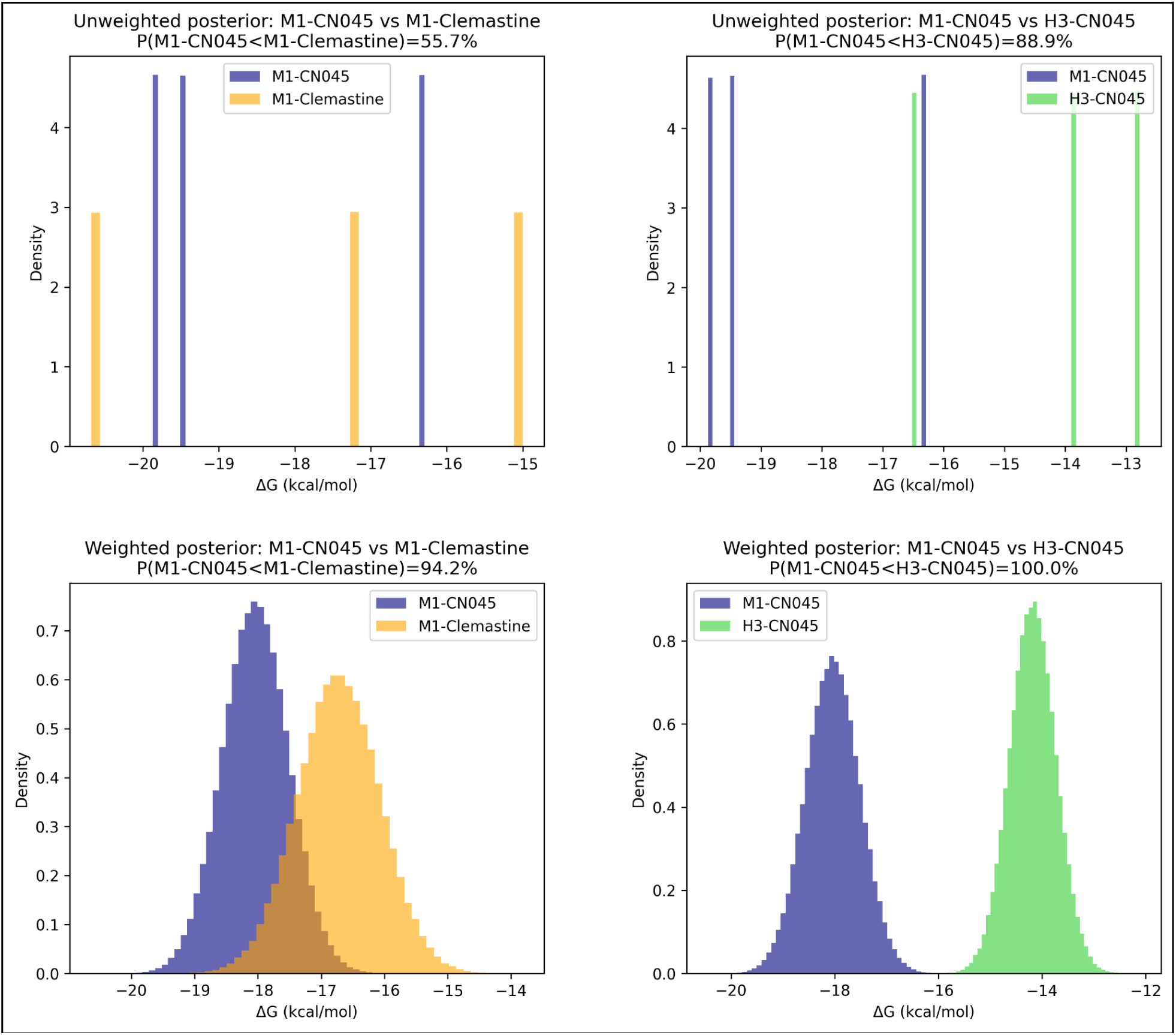
Top – Unweighted posterior overlaps. Posterior distributions of binding free energy (ΔG) for M1-CN045 compared to M1-Clemastine (left) and H3-CN045 (right) using equal-weight replicas. Bottom – Same comparisons as above, but using inverse-variance weighting with per-replica variability estimates (SD(Prop.)). Replicas with smaller SD values contribute more weight to the system’s overall mean. In both cases, probabilities indicate the likelihood that M1-CN045 has a lower (more negative) ΔG than the other systems.

**Table S6:** Dimensions and sizes of the lipid membrane for all 3 protein-ligand complexes.

| System | X Dimension (Å) | Y Dimension (Å) | Total Lipids | Lateral Area (Å<sup>2</sup>) | Area per lipid (Å<sup>2</sup>) | WBOXZ (Å<sup>2</sup>) | WATBOXZ (Å<sup>2</sup>) |
| --- | --- | --- | --- | --- | --- | --- | --- |
| M1-Clemastine | 80.073 | 80.073 | 164 | 6411.6 | 39.1 | 22.5 | 64.296 |
| M1-CN045 | 80.265 | 80.265 | 165 | 6442.5 | 39 | 12.066 | 53.891 |
| H3-CN045 | 80.351 | 80.351 | 167 | 6456.3 | 38.7 | 16.773 | 43.07 |

**Figure S9:**
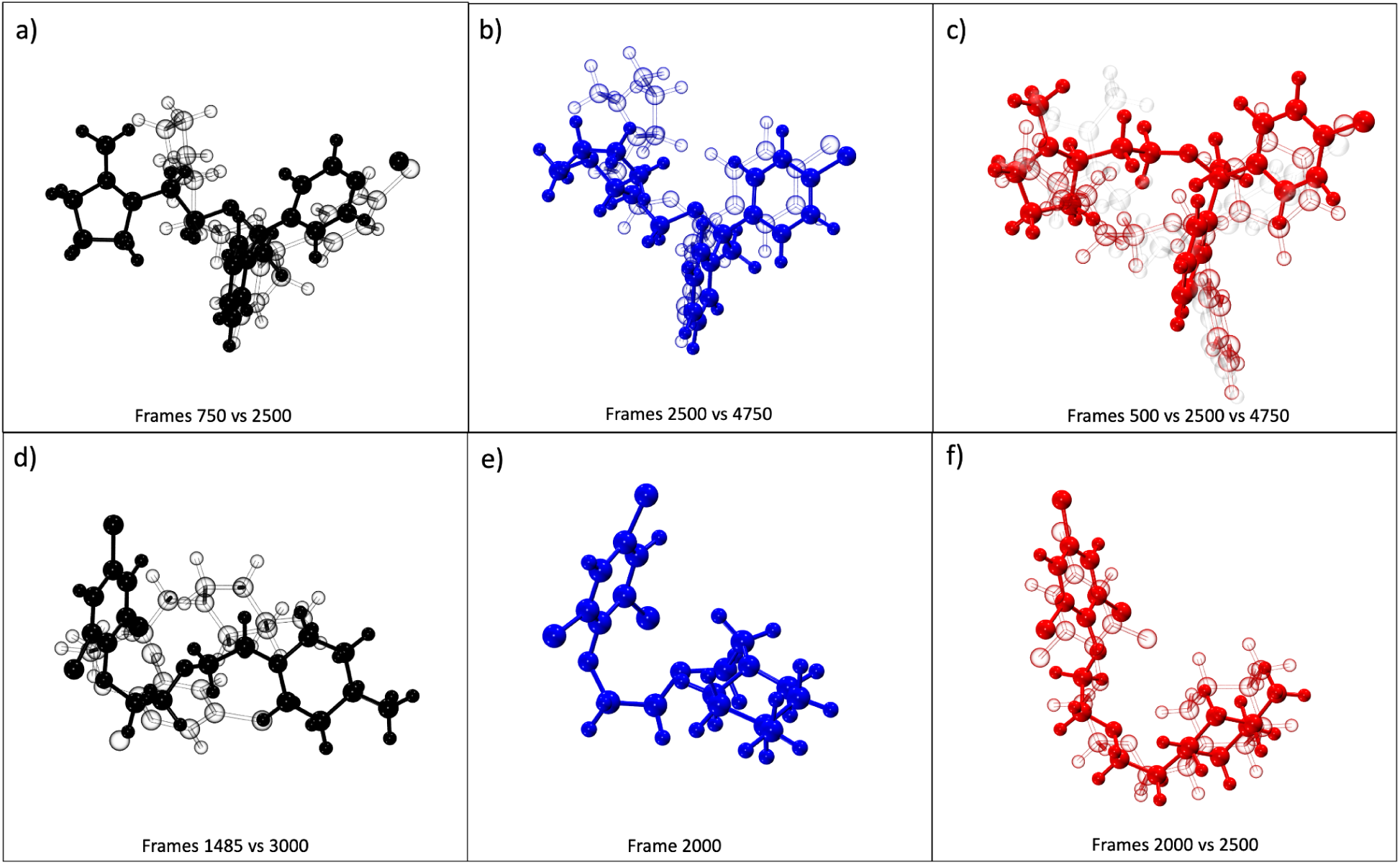
Selected conformations observed in clemastine (top row) and CN045 (bottom row) during the simulations. Structures are colored by replica number (black = replica 1, blue = replica 2, red = replica 3). Each post-processed trajectory contains 5,000 frames corresponding to 1,000 ns. The frames shown here correspond to the nanosecond windows discussed in Sections 3.3.1 and 3.3.2 of the main text. The lower-numbered frame corresponds to the darker structure, whereas the higher-numbered frame corresponds to the transparent or glass-like structure. For clemastine, the predominant conformations appear to be defined by hinging along the aliphatic ether chain, which changes the separation between the aromatic rings and the pyrrolidine group. In addition, approximately 90° rotations of the pyrrolidine group are often observed during these hinging transitions (most clearly in panels a and b). The major change observed for CN045 during replica 1 is characterized by a sudden increase in compactness (panel e) that persists for the remainder of the simulation. Replica 2 remains comparatively constant, with only subtle deviations from the structure seen in frame 2000 (panel f). Replica 3 (panel g) undergoes frequent transitions between two RMSD regions, driven largely by rotations in the piperidine group, similar to what is observed for clemastine.

**Fig. S10:**
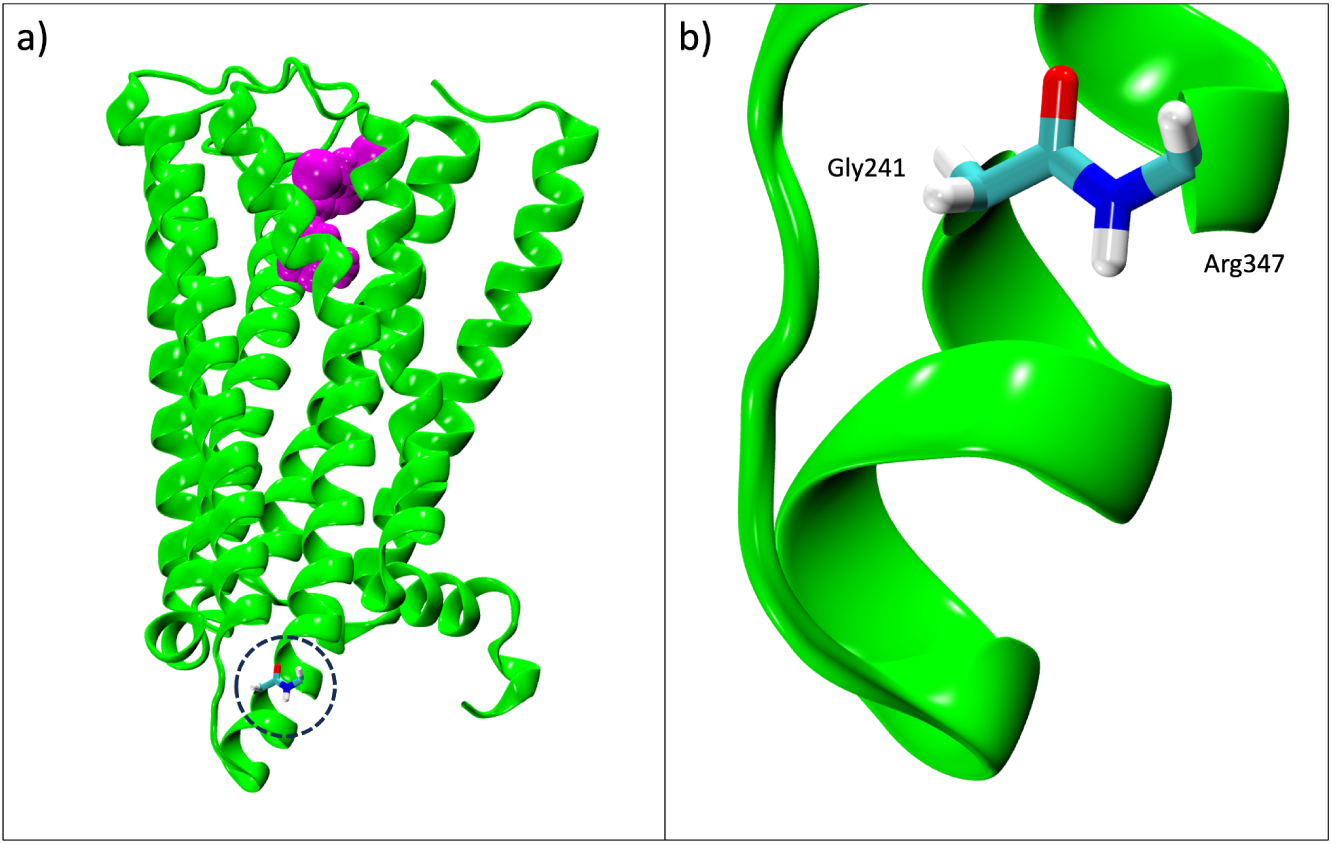
a) The overall structure of the H3 receptor (green) with a bound CN045 molecule (magenta), also highlighting the location of the ICL3 junction. b) A zoomed-in look at the backbone merger between Gly241 and Arg347, creating a continuous backbone across the truncated ICL3 junction.

**Fig. S11:**
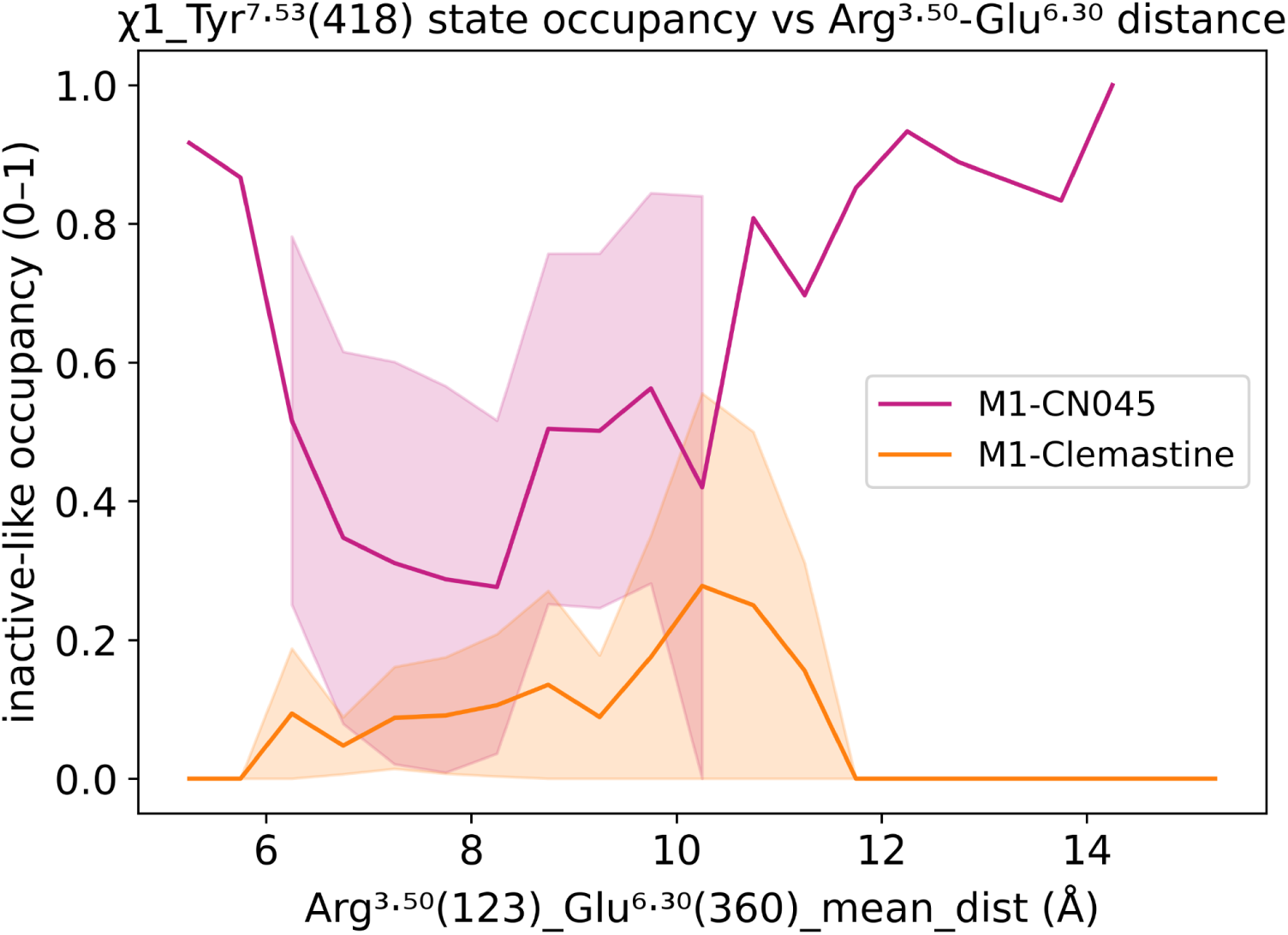
Correlated map of inactive-like occupancy (based on being within a ± 25° threshold of the known inactive state χ1 of +47.5°) and binned distance of the “ionic lock” between Arg^3.50^(123) and Glu^6.30^(360).

